# A quantitative map of nuclear pore assembly reveals two distinct mechanisms

**DOI:** 10.1101/2021.05.17.444137

**Authors:** Shotaro Otsuka, Jeremy O. B. Tempkin, Wanlu Zhang, Antonio Z. Politi, Arina Rybina, M. Julius Hossain, Moritz Kueblbeck, Andrea Callegari, Birgit Koch, Natalia Rosalia Morero, Andrej Sali, Jan Ellenberg

## Abstract

Understanding how the nuclear pore complex (NPC) assembles is of fundamental importance to grasp the mechanisms behind its essential function and understand its role during evolution of eukaryotes^1–4^. While at least two NPC assembly pathways exist, one during exit from mitosis and one during nuclear growth in interphase, we currently lack a quantitative map of the molecular assembly events. Here, we use fluorescence correlation spectroscopy (FCS) calibrated live imaging of endogenously fluorescently-tagged nucleoporins to map the changes in composition and stoichiometry of seven major modules of the human NPC during its assembly in single dividing cells. This systematic quantitative map reveals that the two assembly pathways employ strikingly different molecular mechanisms, inverting the order of addition of two large structural components, the central ring complex and nuclear filaments. The dynamic stoichiometry data underpinned integrative spatiotemporal modeling of the NPC assembly pathway, predicting the structures of postmitotic NPC assembly intermediates.

---

The nuclear pore complex (NPC) is the largest non-polymeric protein complex in eukaryotic cells. It spans the double membrane of the nucleus (nuclear envelope; NE) to mediate the macromolecular transport between the nucleus and the cytoplasm. To achieve this essential function, the NPC forms an octameric proteinaceous channel composed of multiples of eight of over 30 different nucleoporins (Nups) that form 6–8 protein modules, the NPC subcomplexes^1,2^. Therefore, more than 500 individual proteins have to come together to assemble one nuclear pore, which has the mass of tens of ribosomes. NPCs are thought to represent a key step in the evolution of endomembrane compartmentalization that allowed ancestral eukaryotes to separate their genome from the cytoplasm^3,4^.

In proliferating cells, there are two main pathways to assemble the NPCs. During nuclear assembly after mitosis, the NPCs form together with nuclear membranes to rapidly build new nuclei in the daughter cells (called postmitotic NPC assembly). During nuclear growth in interphase, NPCs then continue to assemble continuously for homeostasis (referred to as interphase assembly). Research over the last decade has revealed that postmitotic and interphase NPC assembly possess distinct kinetic, molecular and structural features^5–12^, suggesting that two fundamentally different mechanisms build the same protein complex. In postmitotic assembly, several thousand NPCs assemble within a few minutes during sealing of the initially fenestrated nuclear membranes, whereas interphase NPC assembly occurs more sporadically, requires about one hour, and involves a new discontinuity in the double membrane barrier of the NE. Studies using molecular depletions have shown that the Nup ELYS is required for postmitotic assembly but appears dispensable for interphase assembly^5^, whereas the membrane curvature-sensing domain of Nup133^6^, Pom121 and Sun1^7,8^, the import of Nup153 into the nucleus^9^, and Torsins^10^ seem to be required only for interphase assembly. Recent studies correlating real time imaging with three-dimensional electron microscopy have revealed that postmitotic NPC assembly proceeds by radial dilation of small membrane openings^11^, while in interphase, assembly induces an asymmetric inside-out fusion of the inner and outer nuclear membranes^12^.

However, how several hundred proteins self-organize to form the NPC channel *via* these two distinct assembly pathways has remained largely enigmatic. It is technically challenging to locate the transient and rare assembly events, which has prevented investigation of the structure of assembly intermediates by either cryo-EM tomography or super-resolution microscopy. In addition, the large number of building blocks and their cooperativity often leads to complex nonlinear kinetics that can only be interpreted mechanistically using computational modeling of the structures formed during assembly. While we have some information about the dynamic addition of Nups after mitosis^13,14^, only sparse dynamic data is available for interphase assembly^15,16^. Importantly, these earlier studies could not distinguish postmitotic and interphase assemblies, whose co-occurrence in different regions of the nucleus was only discovered later; moreover, these studies provided only qualitative descriptions as they relied on ectopic expression of fluorescently-tagged Nups. Kinetic data about NPC assembly that can distinguish between the postmitotic and interphase pathways is required, including the copy numbers of Nups that assemble into forming NPCs over time. Such data would enable modeling of the assembly process and allow us to understand the two assembly mechanisms.

## Quantitative imaging of ten nucleoporins

To quantitatively analyze the changes in concentration of Nups at the NE during exit from mitosis and nuclear growth in G1, we genome-edited HeLa cells, homozygously tagging the endogenous genes for different Nups with mEGFP or mCherry. Building previous work^11,12,17,18^, we created a set of ten endogenously tagged Nups that systematically represent the major building blocks of the fully assembled pore, including the nuclear filament protein Tpr, the nuclear Nup153, the Y-complex members Nup107 and Seh1, the central ring complex members Nup93 and Nup205, the central channel protein Nup62, the transmembrane protein Pom121 as well as the cytoplasmic filament proteins Nup214 and Nup358. Homozygous tagging was verified by careful quality control of the genome edited monoclonal cell lines^19^, ensuring that the tagged subunit was expressed at physiological levels, localized to the NPC and that cell viability and mitotic progression were normal (Figs. 1 and 2). The fusion proteins are likely functional, given that most Nups show strong phenotypes upon knock-out or depletion^20^.

**Fig. 1.**
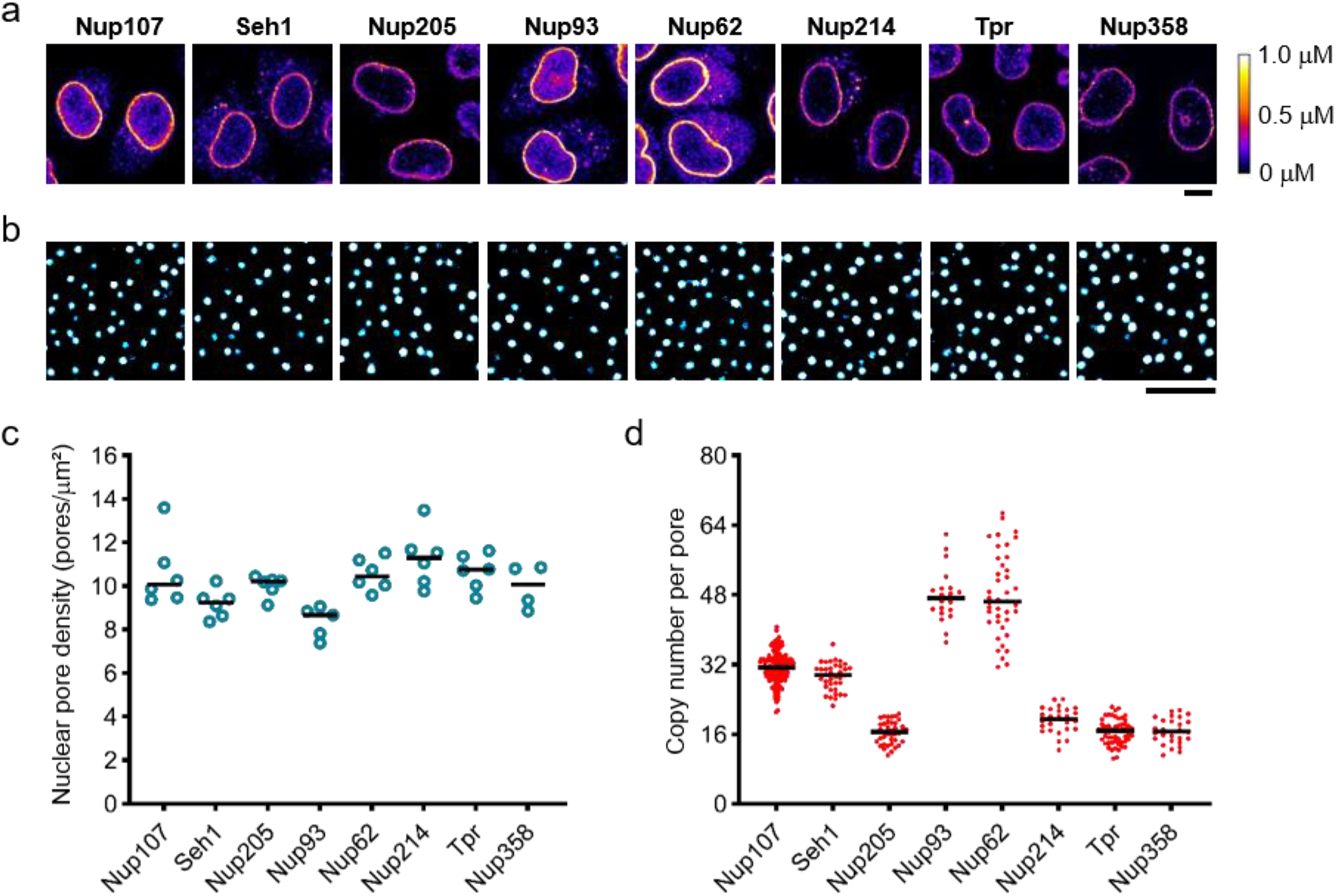
Quantitative imaging of GFP-knock-in nucleoporin (Nup) cell lines. **a**, Genome-edited HeLa cells with homozygously mEGFP-tagged Nups observed by confocal microscopy. Fluorescent intensity was converted into protein concentration by FCS-calibrated imaging^22^. Images were filtered with a median filter (kernel size: 0.25 × 0.25 μm) for presentation purposes. Scale bar, 10 μm. **b**, **c**, Stimulated emission depletion (STED) microscopy. The genome-edited cells were stained with anti-Nup62 antibody and imaged (**b**), and then the density of nuclear pores was quantified (**c**). The plot is from 6, 6, 6, 5, 6, 6, 6, and 4 cells for Nup107, Seh1, Nup205, Nup93, Nup62, Nup214, Tpr and Nup358, respectively. Scale bar, 1 μm. **d**, Calculated copy number of Nups per nuclear pore. The plot is from 241, 37, 41, 20, 41, 26, 55, and 28 cells for Nup107, Seh1, Nup205, Nup93, Nup62, Nup214, Tpr and Nup358, respectively. The median is depicted as a line.

**Fig. 2.**
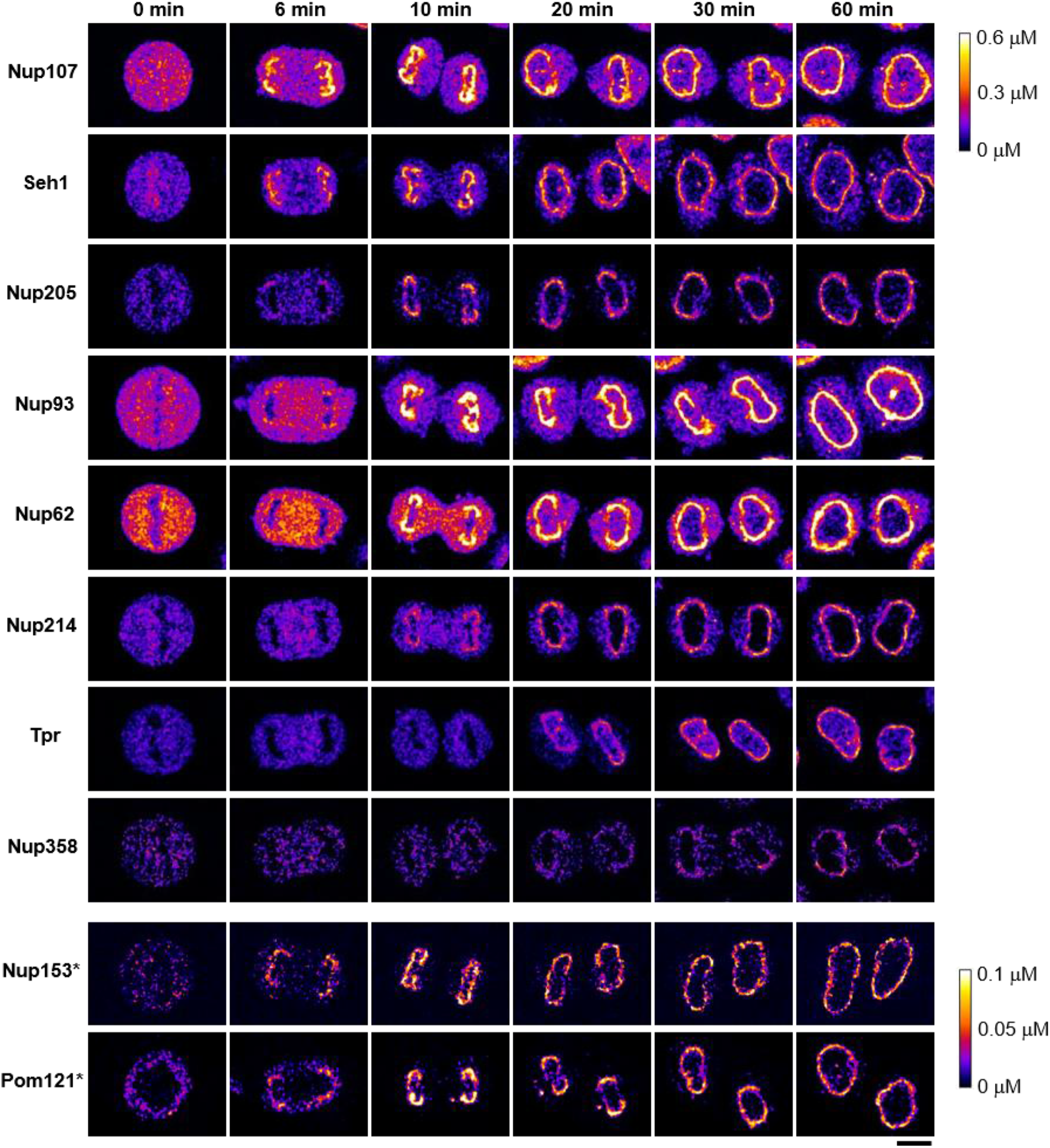
Dynamic concentration maps of Nups after anaphase onset (AO). HeLa cells whose Nups are endogenously tagged with mEGFP or mCherry were imaged every 30 sec by three-dimensional (3D) confocal microscopy. Single confocal sections are shown. Images were calibrated by FCS to convert fluorescence intensities into cellular protein concentration. *For Nup153 and Pom121, these Nups are not fully validated to be homozygously-tagged. Images were filtered with a median filter (kernel size: 0.25 × 0.25 μm). Scale bar, 10 μm.

To characterize the fluorescently-tagged Nups, we first performed super-resolution (STED) microscopy to determine NPC density in fully grown nuclei of the knock-in cell lines, showing that homozygous tagging had little effect on NPC density that was comparable within 15% between all cell lines with an average of 10.1 NPC per μm^2^ (Fig. 1b, c), in good agreement with our previous estimates by electron microscopy of HeLa cells^12^. We then used fluorescence correlation spectroscopy (FCS) calibrated confocal microscopy^21,22^ to determine the concentration and total number of the Nups at the NE in living cells (Extended Data Table 1, for details see Methods). Using the measured NPC density and the Nup concentration at the NE, we could calculate the average copy number of each Nup per NPC (Fig. 1d). This data showed that the investigated Nups were on average present in 16, 32, or 48 copies per pore, as expected from the eightfold symmetry of the complex and overall in good agreement with previous estimates by mass spectrometry^23^.

## Dynamic change of nucleoporin numbers

We then used our validated cell line resource to quantitatively image the Nups during both postmitotic and interphase NPC assembly, from metaphase until the end of the rapid nuclear growth phase in G1, two hours after anaphase onset. To this end, we performed systematic FCS-calibrated 3D confocal time-lapse microscopy^17^ (Fig. 2). Using the single molecule fluctuation calibration, the 4D imaging data could be converted into maps of subcellular protein concentration (Fig. 2). Counterstaining live nuclei with SiR-DNA^26^, enabled computational image segmentation to measure the soluble cytoplasmic pool and the NE associated pool over time^17^. Temporal alignment to anaphase onset then allowed us to compare the dynamic association of all Nups with the NE over time (Fig. 2). Overall, the investigated Nups are present in 250,000 to 1,200,000 copies per human metaphase cell. After mitosis, this building material is split between the daughter cells with little detectable new protein synthesis in the first hour after anaphase onset (Extended Data Fig. 2). Notably, 34–53% of the soluble pool present in the cytoplasm in metaphase was rapidly re-localized to the NE during the first hour after exit from mitosis (43, 38, 47, 37, 41, 44, 53, and 34% for Nup107, Seh1, Nups205, 93, 62, 214, Tpr and Nup358, respectively) (Extended Data Fig. 2), indicating that NPC assembly initially relies almost entirely on the pool of building blocks inherited from the mother cell to form the first 4,000–5,000 NPCs^11,12^. We confirmed that the observed GFP-tagged Nups are indeed recruited specifically to NPCs and not in general to the NE surface by single NPC imaging using STED microscopy (Extended Data Fig. 3a,b).

## Quantification of assembly kinetics

Postmitotic and interphase NPC assembly can be observed in the same living cell in different regions of the NE within the first two hours after mitosis^12^. While postmitotic assembly dominates the peripheral “non-core” regions of the NE, the central “inner core” area is only populated with NPCs after exit from mitosis^27^ when dense spindle microtubules have been removed from the DNA surface^28^. Using computational segmentation and assignment of the inner-core and non-core regions (Extended Data Fig. 4a, for detail see Methods modified from^12^), we measured the concentration changes of the ten Nups in these two regions separately. A two-component model of a fast (postmitotic) and a slow (interphase) assembly process fits the experimental data well, allowing us to kinetically unmix the two assembly processes for each Nup (Extended Data Figs. 4b–d and 5). In this way, we could for the first time perform an integrated analysis of the real time kinetics of absolute concentration changes of Nups in all major NPC modules during the two assembly processes at the NE (Fig. 3a). This analysis immediately revealed that the overall duration of the two processes is very different, with postmitotic assembly essentially complete 15 min after anaphase onset, while interphase assembly only reaches a plateau after 100 min, consistent with our previous estimates based on live cell imaging^13,16^ and correlative electron microscopy^11,12^. Both processes reached the same final ratios between the different Nups and thus presumably formed identical NPCs. However, the temporal order in which components were added was distinct, including, for example, an earlier assembly of Pom121 relative to the Y-shaped complex during interphase assembly, consistent with our previous observations^16^.

**Fig. 3.**
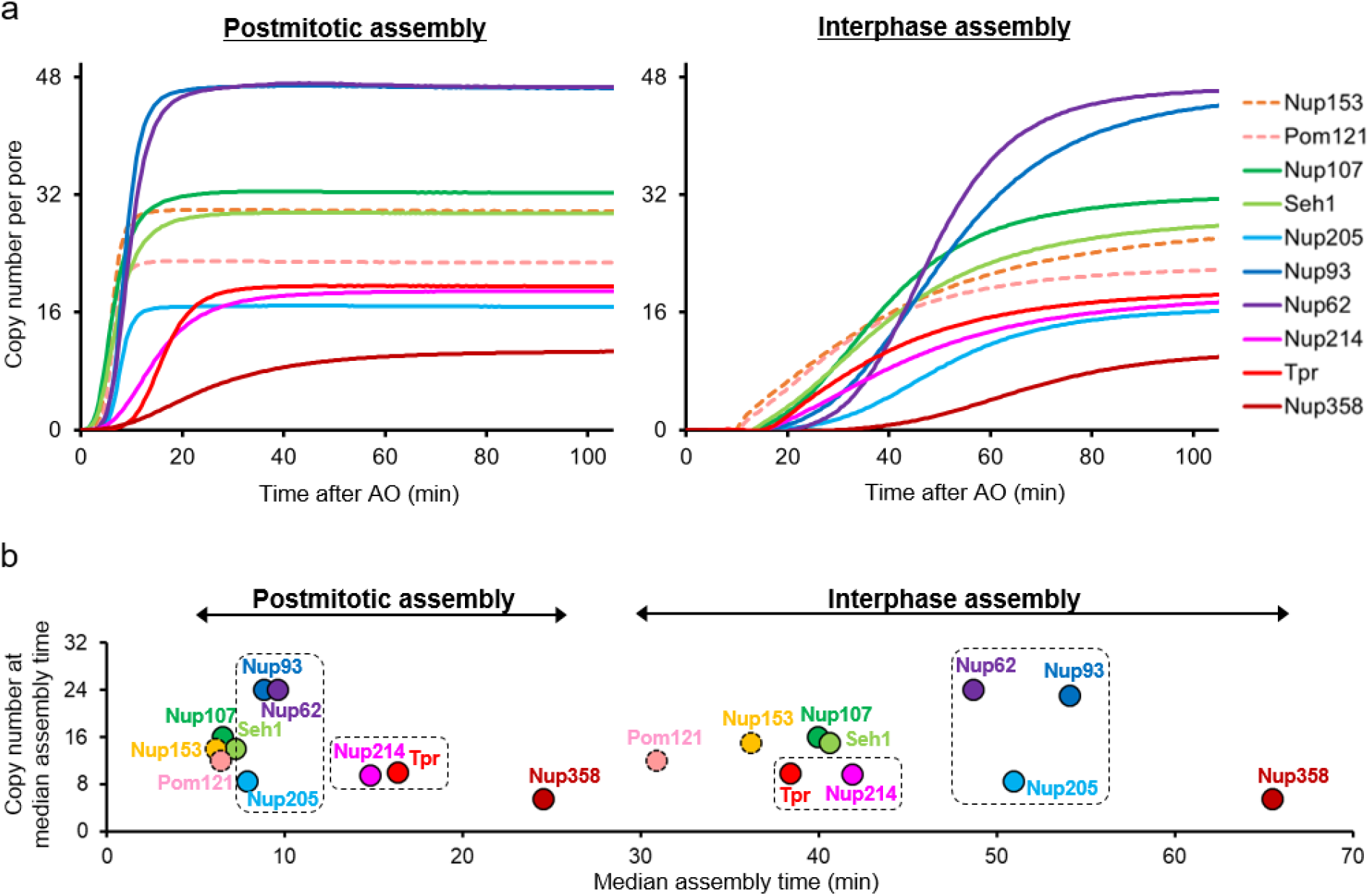
The molecular assembly order and maturation kinetics are distinct for postmitotic and interphase assembly. **a**, Plots of the average copy number per nuclear pore computed from mathematical modeling for postmitotic (left) and interphase (right) assembly (see Methods and Extended Data Figs. 4 and 5 for details). For Nup153 and Pom121 (dashed lines), their absolute amount was estimated using the copy number determined from the previous study (32 for Nup153 and 16 for Pom121)^23^. **b**, The average copy number of individual Nups per nuclear pore are plotted along their median assembly time in postmitotic and interphase assembly pathways. The boxes highlight the Nups that show striking difference in their order of assembly in the two pathways.

To comprehensively investigate the molecular differences between the two assembly processes, we relied on the constant NPC density and changes in nuclear surface area^12^ to convert the NE concentrations of all investigated Nups into their average copy number per assembling NPC over time (Fig. 3a). This result in turn enabled us to estimate changes in subunit stoichiometry of the complex during its assembly in living cells. To facilitate the comparative analysis of the assembly kinetics between the two pathways, in which ten components assemble with different speed and order, we first reduced the dimensionality of the kinetic data. To this end, we assigned a single characteristic time-point of assembly to each Nup, by sigmoidal fitting of its full kinetic signature (Extended Data Fig. 6a). Plotting the copy number *vs* the median assembly time provides an overview of the major molecular differences between the two assembly pathways (Fig. 3b).

While the Y-complex, Pom121, and Nup153 form a core of the first modules that assemble almost simultaneously within one minute in postmitotic assembly, these components are stretched out into a clear temporal order of first Nup153, second Pom121, and third the Y-complex (notably with its two investigated subunits also assembling simultaneously in interphase) over more than ten minutes in interphase assembly. The end of assembly on the other hand is marked for both processes by the addition of the large cytoplasmic filament protein Nup358. The major difference was thus neither in initiation nor termination of assembly, but rather in the middle of the two assembly pathways. During postmitotic assembly, the Y-complex is rapidly combined with components of the central ring, building the inner core of the pore within only three minutes prior to addition of either cytoplasmic or nuclear filament proteins, which follow later. In contrast, during interphase assembly, the Y-complex is first combined with the nuclear filament protein Tpr and the base of the cytoplasmic filament Nup214, while the central ring complex is added later. This observation clearly shows that postmitotic and interphase NPC assembly not only proceed with different speed but also follow a different molecular mechanism using an inverted molecular order between the central ring and nuclear filaments (Fig. 3b, Extended Data Fig. 6b,c).

To validate if Tpr indeed assembles earlier in the interphase assembly pathway, we immuno-stained assembling NPCs by antibodies against Tpr and a later assembling nucleoporin Nup62 and visualised them by 3D-STED super-resolution microscopy. This single NPC imaging demonstrated that a large number of assembling NPCs in the inner-core region in early G1 contain Tpr but not Nup62, while most of the assembled NPCs in the non-core region of the same cells contain both Tpr and Nup62 (Fig. 4a,c), confirming the surprisingly early recruitment of Tpr in interphase assembly pathway. In addition to cells in early G1, we also examined cells later in interphase during S- and G2-phase. Consistently, we found that a significant number of NPCs also contain Tpr but not Nup62 (Fig. 4b,c), indicating that interphase assembly at later cell-cycle stages follows the same order.

**Fig. 4.**
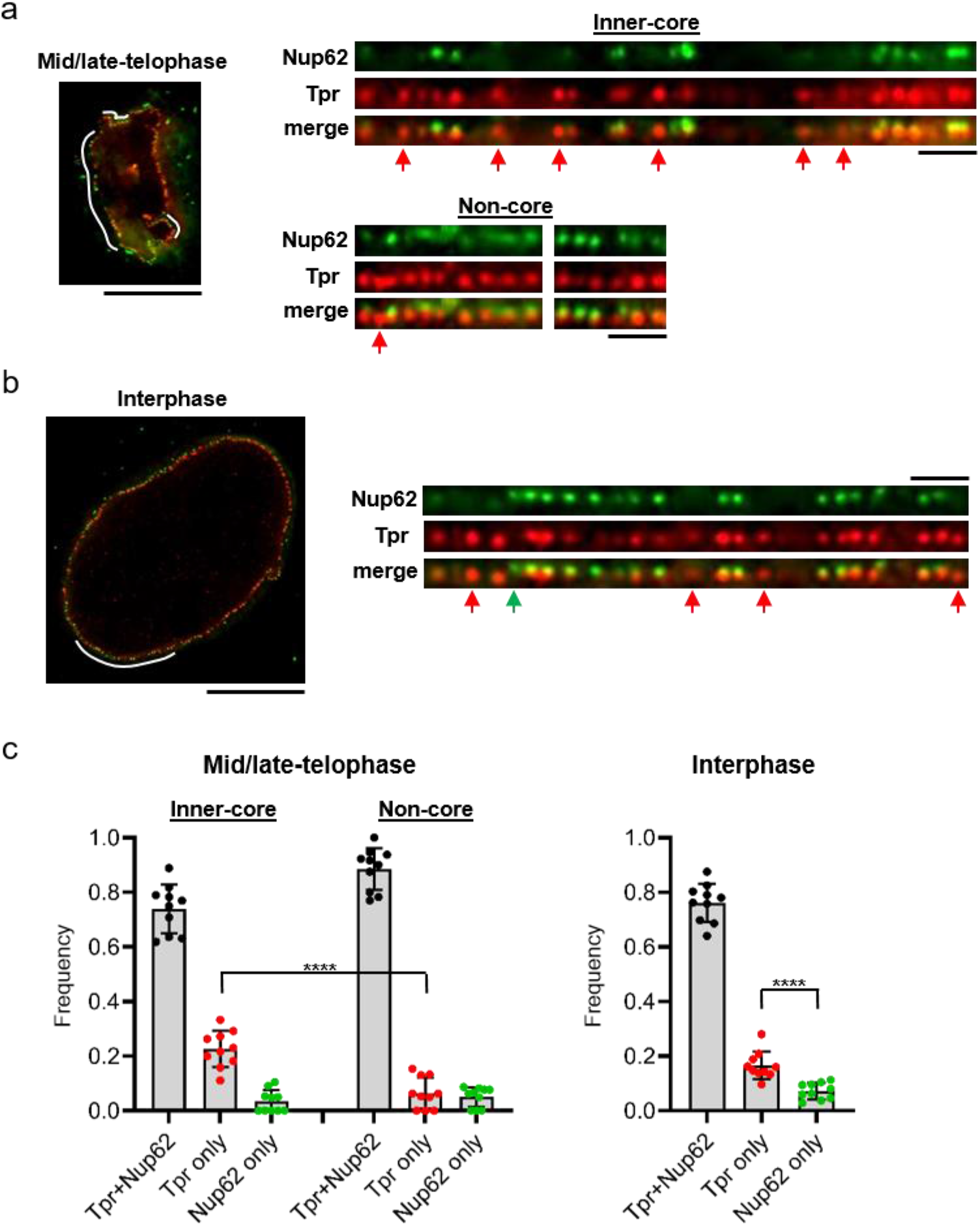
Single nuclear pore observation by super-resolution microcopy confirmed the earlier recruitment of Tpr than Nup62 in interphase assembly pathway. **a,b**, 3D-STED imaging of Nup62-mEGFP genome-edited cells that were stained with a GFP-Nanobody and an anti-Tpr antibody at different cell-cycle stages (**a**: mid/late telophase; **b**: interphase). Scale bars in the left and right panels, 10 μm and 1 μm. The regions of the nuclear envelope indicated by white lines in the left images are flattened and shown in the right panels. The nuclear pores that contain only Tpr or Nup62 are indicated by red and green arrows, respectively. **c**, The frequency of the nuclear pores that contain both Tpr and Nup62, only Tpr and only Nup62 at each cell-cycle stage. The data are from 10 cells for each stage. Error bars represent the s.d. of the mean. ****P=0.000017 and P=0.000080 for mid/late telophase and interphase, respectively; unpaired two-tailed t-tests.

## Integrative modeling of the assembly

To obtain a more comprehensive mechanistic view, we computed a spatiotemporal model of the macromolecular assembly pathway, based on our dynamic multimolecular stoichiometry data in combination with the available ultrastructural data about NPC assembly^11^ and the partial pseudoatomic model of the mature human NPC^29,30^. We modeled here only the postmitotic assembly, as it showed the kinetic hallmarks of a sequential process and is known to proceed by dilating an existing membrane pore with a smoothly growing proteinaceous density^11^. We focused on the Nups contained in the structural model of the human NPC, including Nup107 and Seh1 for the Y-complex as well as Nup93, Nup205, and Nup62 for the central ring/channel complex^29,30^. We constrained their copy numbers by our stoichiometry data for the time points for which correlative electron tomography data is available^11^, allowing us to use the membrane shapes and associated protein densities from tomography to model the Nup configurations at these time points. To compute a spatiotemporal assembly pathway model, we generalized our integrative modeling method for determining static structures of macromolecular assemblies^31,32^. In outline (see Methods for details), we first compute ensembles of Nup configurations at discrete time points, connect pairs of these configurations at adjacent time points into assembly trajectories, and finally rank the alternative trajectories by fit to our data.

We focus on the best fitting macromolecular assembly pathway that accounts for over 85% of the posterior model likelihood (the second-scoring model accounts for only 14.5%, Fig. 5, Extended Data Fig. 7). This trajectory starts by formation of a single nuclear ring, composed of eight Y-complexes, concomitantly with an initial accumulation of the FG-repeat protein Nup62 and the inner-ring complexes (the Nup93-Nup188-Nup155 and Nup205-Nup93-Nup155 complexes and Nup155) in the center of the membrane hole (Fig. 5; 5 and 6 min). We experimentally validated that Nup155 and Nup188 assemble as the model predicted (Extended Data Figs. 8, 9, Supplementary Fig. 1). The cytoplasmic Y-complex is then added on the cytoplasmic side, before the second set of the Y-complex ring assembles on the nuclear side, again one eight-membered ring after another (Fig. 5; 8 and 10 min). In the center of the pore, Nup62 dilates from an amorphous mass into a small ring and associates with inner-ring Nups to form the central ring complex. This “nuclear-ring-first” assembly mechanism is consistent with the observation of an eight-fold symmetric protein density on the inner nuclear membrane at early stages of assembly^11^.

**Fig. 5.**
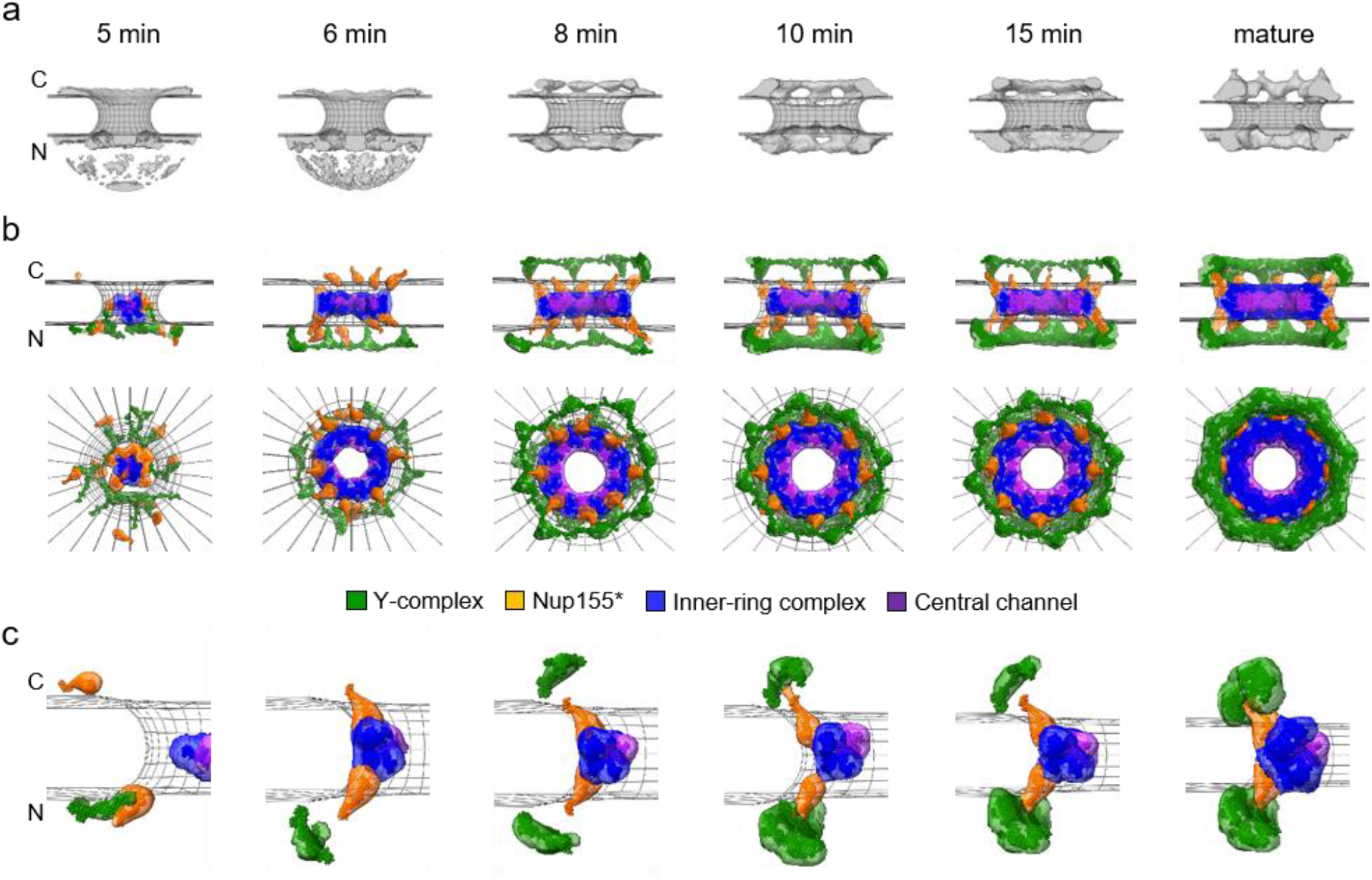
Integrative model of the postmitotic NPC assembly pathway. **a**, Protein density (grey) overlaid with the NE surface (wireframe model, grey) at each time point^12^ used for integrative modeling. C, cytoplasm; N, nucleoplasm. The dome-like density in the nucleoplasmic side at 5 and 6 min is the noise from electron tomography. **b**, The best-scoring model of the postmitotic assembly pathway (top and side views). The uncertainty of each Nup localization is indicated by the density of the corresponding color: Y-complex (green), *isolated fraction of Nup155 that is not forming a complex with Nup93, Nup188, and Nup205 (orange), the Nup93-Nup188-Nup155 and Nup205-Nup93-Nup155 complexes (blue), and the Nup62-Nup58-Nup54 complex (purple). **c**, The enlargement of one of the spokes. The model has a much higher score than all the other lower-scoring pathways (Extended Data Fig. 7). The pseudoatomic model of the native NPC structure^29,30^ we imposed as the endpoint of the assembly pathway does not include the Y-complex-bound fraction of Nup205 nor Nup214-bound fraction of Nup62, thus containing only 16 of the 40 copies of Nup205^24,25^ and 32 of the 48 copies of Nup62^36^ in the fully mature NPC; how the remaining 24 copies of Nup205 and 16 copies of Nup62 are assembled remains elusive.

## Discussion

Our data revealed that the two assembly pathways are strikingly different. Although it had been shown that postmitotic and interphase assembly pathways have different molecular requirements^5–12^, very little was known about the interphase pathway due to its rare and sporadic nature. While relative kinetics were available for some Nups^15,16^, how key Nups composing cytoplasmic filaments, central channel, and nuclear basket, or multiple subunits of the Y-shaped and inner-ring complexes assemble in interphase had not been studied. Here, we provide the first measurements on subunit composition and stoichiometry for both postmitotic and interphase assembly for ten Nups that represent all major building blocks of the NPC. Our data thus systematically reveals the main molecular differences between the two assembly pathways. During postmitotic assembly, the Y-complex is rapidly combined with components of the central ring, building the inner, and at this time already transport competent^33^, core of the pore prior to addition of either cytoplasmic or nuclear filament proteins. By contrast, during interphase assembly, the Y-complex is first combined with the nuclear filament protein Tpr and the base of the cytoplasmic filament Nup214, while the central ring complex is added much later.

The molecular pathway we observed for interphase assembly makes new predictions about its unique, inside out evaginating, mechanism. The cell first not only builds the two nuclear Y-rings but also accumulates the material for their cytoplasmic counterparts and then combines them with the nuclear filament proteins, all prior to the time of membrane fusion^12^. This mechanism suggests that the cytoplasmic Y-rings, including the base of the cytoplasmic filament Nup214, are already “prebuilt” within the inner membrane evagination, where the small available volume would predict that they must be present in a very different structure than in the fully mature pore after fusion. Surprisingly, the central ring complex, which is a core structural element between the nuclear and cytoplasmic Y-rings in the mature pore is added later, suggesting that during interphase assembly, nuclear transport control is added after the membrane fusion step. The unexpectedly early presence of Tpr suggests, although does not prove, a potential role of Tpr in the initiation of interphase assembly. On the one hand, this long coiled-coil multimeric protein component could be involved bending the inner nuclear membrane to create the invagination, potentially providing force by unfolding or restructuring a coiled-coil bundle^1,2^. On the other hand, considering previous reports that Tpr together with the kinase ERK is required to control the spacing between NPCs^34^, Tpr-based signaling might also be important in assembly site selection, to prevent simultaneous assembly of multiple NPCs in close proximity with each other, which would likely result in abnormal distortions of the inner nuclear membrane.

Our data on interphase assembly in human cells furthermore revealed striking differences to budding yeast that undergoes a closed mitosis and lacks the postmitotic pathway^35^; (i) the Tpr homologues Mlp1/2 assembles late in yeast, (ii) human cells exhibit a synchronous assembly of Y-complex components, while in yeast the stem (e.g. Nup107) and the head (e.g. Seh1) of the Y-complex do not assemble simultaneously, and (iii) the central ring components assemble after the Y-complex components in human cells, whereas they assemble together in yeast^35^. These differences could be due to the fundamental differences in structure (e.g., presence of a nuclear lamina) and/or cell cycle remodeling of the NE (closed *vs* open mitosis) between human and yeast cells.

Our dynamic stoichiometry data, combined with previous electron tomography data on membrane topology and protein volume, allowed us for the first time to predict the structures of the NPC assembly intermediates using integrative spatiotemporal modeling. We focused on the topologically simpler postmitotic assembly. The resulting model allows to make multiple new mechanistic predictions regarding postmitotic assembly that will help guide future work. For example, the model suggests that the central ring complex might be needed to prevent sealing of ER holes after mitosis, and that the hydrophobic FG-repeats of central channel Nups might play a role in the dilation of the small membrane hole into the larger, NPC sized, channel. Our pathway model was constrained by the fully assembled NPC structure^29,30^ containing 16 copies of Nup205 and 32 copies of Nup62; thus, it does not address how additional copies of Nup205 and Nup62 that were reported in recent structural studies and a proximity-dependent biotin identification study^24,25,36^ may be assembled.

Our combined experimental and computational approach for determining macromolecular assembly pathways is likely to benefit from additional experimental data provided by emerging methods, such as 3D super-resolution microscopy for imaging the molecular architecture of the NPC intermediates^37,38^ and dynamic super-resolution methods for mapping structural changes in real time^39^. We expect that this additional information will allow higher resolution modelling of even more complex assembly mechanisms, such as the interphase NPC assembly that involves much more dramatic membrane topology and protein conformation changes.

The need for better methods to determine dynamic and variable copy numbers of Nups reliably at the level of single protein complexes under physiological conditions is further highlighted by the fact that our live cell-based approach resulted in copy number estimates that differ from estimates derived by other approaches for four out of the eleven investigated Nups. The main methods employed in the field, including quantitative mass spectrometry from cell populations^23^, fitting of single protein structures into highly averaged cryo-EM densities^24,25^, as well as calibrated imaging of fluorescently-tagged knock-in proteins in single living cells as employed in our study, currently all have limitations in this regard. For example, cryo-EM based averaging often selects “complete” NPCs, to obtain a “fully occupied” average NPC model, while live cell imaging data averages NPC stoichiometry from all pores in one cell and thus includes pore-to-pore variability, which can result in lower estimates. On the other hand, while homozygous genome editing of a fluorescent tag ensures close to complete labelling and biological function of the fusion protein, it can still affect the expression level, which could lead to a different occupancy within the NPC.

Four of the eleven investigated Nups are relevant to discuss in the light of these methodological limitations, i.e. Nup153, Pom121, Nup205, and Nup358. For the first two, our estimates were much lower than expected from the structural model of the fully assembled state we imposed as the end-state of our pathway model (Fig. 2, Extended Data Fig. 1). We therefore regarded them as effectively subphysiologically expressed and normalized their stoichiometry to the expected number of copies in the mature pore for comparison with other Nups (Fig. 3a). For Nup205 and Nup358, our copy number estimates were 40-50% lower than expected, even though for Nup205 our cell lines express physiological levels of the fusion protein, while for Nup358 this correlates to a reduced level of expression (Supplementary Fig. 2). More work is needed to reliably measure the dynamic and variable composition of the NPC, especially during its assembly, and incorporate this data into modeling.

Beyond the assembly mechanism, the two pathways we have mapped here start to shed light on the role of the NPC during evolution of the endomembrane system in eukaryotes^3,4^. We speculate that the modern NPC combines ancient membrane bending (e.g., coiled-coil filaments, such as Tpr) and membrane hole plugging/dilation (e.g., central ring complex and FG-repeat proteins) modules that were potentially previously used separately for extruding the cell surface or keeping transport channels open in the endomembranes around the genome. The key evolutionary innovation might lie in combining and controlling these activities, potentially by the eight-membered ring architecture of the nuclear Y-complex. In the future, comparing NPC assembly pathways in different species on the eukaryotic evolutionary tree might help us to understand how the assembly of complex modern protein machines reflects their evolutionary origins.

## Methods

### Cell culture

Wildtype HeLa kyoto cells (RRID: CVCL_1922) were kind gift from Prof. Narumiya in Kyoto University, and the genome was sequenced previously^40^. Cells were grown in high glucose Dulbecco’s Modified Eagle’s Medium (DMEM) containing 4.5 g/l D-glucose (Sigma Aldrich, St. Louis, MO) supplemented with 10% fetal calf serum (FCS), 2 mM l-glutamine, 1 mM sodium pyruvate, and 100 μg/ml penicillin and streptomycin at 37 °C and 5% CO_2_. The mycoplasma contamination was inspected by PCR every 2 or 3 months and was always negative.

### Genome editing

Monomeric enhanced GFP (mEGFP) and mCherry were inserted into the genome using zinc finger nucleases, CRISPR-Cas9 nickases, or Alt-R® S.p. HiFi Cas9 Nuclease V3. The following six cell lines had been generated and published previously: Nup62-mEGFP^18^, mEGFP-Nup107^12^, mEGFP-Nup205^11^, mEGFP-Nup214, mEGFP-Nup358 (also called RanBP2) and Tpr-mEGFP^17^. The following five cell lines were generated in this study: mEGFP-Seh1, Nup93-mEGFP, mEGFP-Nup153, Nup188-mEGFP and Pom121-mCherry. The gRNA sequences used for generating these cell lines are summarized in Supplementary Table 1. For the Nup93-mEGFP cell line, CRISPR-Cas9 nickases and the donor plasmid were transfected by electroporation (Neon Transfection System, Thermo Fisher Scientific, Waltham, MA) instead of a polymer-mediated transfection reagent.

### Western blot

Cells were lysed on ice in RIPA buffer (50 mM Tris-HCl, 150 mM NaCl, 1.0% Triton X-100, Sodium Deoxycholate 1%, SDS 0.1%, EDTA 2 mM), pH 7.5, supplemented with complete EDTA-free protease inhibitor cocktail (Sigma Aldrich), PhosSTOP (Sigma Aldrich), and 0.1 mM Phenylmethylsulfonyl fluoride (Sigma Aldrich). The cell lysates were snap-freezed in liquid Nitrogen and quickly thawed at 37 °C two times. The lysates were then centrifuged at 16,000 g at 4°C for 10 min and the supernatant was used for immunoblot analysis. Protein concentration was quantitated using the Pierce BCA Protein Assay Kit (Thermo Fisher Scientific). For Nup188 (Extended Fig. 9b), the cell lysates were run onto NuPAGE®4–12% Bis-Tris Gels (Novex Life Technologies, Waltham, MA) and transferred onto PVDF membrane using the Bio-Rad transfer system. After blocking with 4% milk solution (nonfat milk powder in PBS + 0.05% Tween 20), the proteins were labelled by anti-Nup188 (Catalog No. A302-322A, Bethyl Laboratories, Montgomery, TX, 1:5000) and anti-γ-Tubulin (Cat. No. T5192, Sigma Aldrich, 1:1000) antibodies. Subsequently anti-rabbit IgG horseradish peroxidase (HRP)-conjugated secondary antibody (Cat. No. W4011, Promega, Madison, WI) was used to detect the protein of interest with chemiluminescence reaction. For Nup205 and Nup358 (Supplementary Fig. 2), the lysates were run onto NuPAGE®3–8% Tris-acetate gels (Novex Life Technologies), and anti-Nup205 (Catalog No. ab157090, Abcam, Cambridge, UK, 1:1000), anti-Nup358 (Catalog No. HPA023960, The Human Protein Atlas, 1:1000), and anti-Vinculin (Cat. No. ab219649, Abcam, 1:1500) antibodies were used to detect the proteins. Simple western assays were also performed in a Jess instrument (ProteinSimple, Santa Clara, CA) using 66–440 kDa capillary cartridges in accordance with the provider’s instructions. The anti-Nup205, Nu358, and γ-Tubulin antibodies were used at 1:50 dilution. For Nup205, mouse anti-γ-Tubulin antibody (Catalog No. T5326, Sigma) was used instead of the rabbit anti-γ-Tubulin antibody to adjust the signal intensity. The secondary antibodies used were the ones provided by the company at a ready-to-use dilution (goat anti-mouse HPR-conjugated secondary antibody, Catalog No. 040-655, Bio-Techne, Minneapolis, MN; goat anti-rabbit HPR-conjugated secondary antibody, Catalog No. 040-656, Bio-Techne).

### FCS-calibrated live-cell imaging and estimation of Nup copy numbers per NPC

All the live imaging was performed in the following way. (i) Wild-type cells, (ii) wild-type cells transfected with mEGFP using Fugene6 (Promega), (iii) mEGFP-Nup107 genome-edited cells, and (iv) the cells of another mEGFP-Nup genome-edited cell line, were seeded on each well of 8-well Lab-Tek Chambered Coverglass (Thermo Fisher Scientific). On the day of live-cell imaging, DMEM was replaced by imaging medium: CO_2_-independent medium without phenol red (Invitrogen) containing 20% FCS, 2 mM l-glutamine, and 100 μg/ml penicillin and streptomycin. The imaging medium was supplemented with 50 nM silicon–rhodamine (SiR) Hoechst to stain DNA^26^. Cells were incubated inside the microscope-body-enclosing incubator at 37 °C for at least 30 min before imaging. For Nup188-mEGFP genome-edited cells, the following medium was also used instead of the imaging medium: 30 mM HEPES pH 7.4 containing 9.3 g/l Minimum Essential Medium Eagle (Sigma Aldrich), 10% FCS, 1% MEM Non-Essential Amino Acids (Thermo Fisher Scientific/Gibco), and 100 μg/ml penicillin and streptomycin.

Calibrated imaging using fluorescence correlation spectroscopy (FCS) was carried out as described in a previous report^22^, using Fluctuation Analyzer 4G 150223 (https://www-ellenberg.embl.de/resources/data-analysis), FCSFitM v0.8 (https://git.embl.de/grp-ellenberg/FCSAnalyze), FCSCalibration v0.4.2 (https://git.embl.de/grp-ellenberg/FCSAnalyze), RStudio 1.1.383, R 3.4.1, and Python v3.6.8. Briefly, the confocal volume was determined by performing FCS using a dye with known diffusion coefficient and concentration (Alexa Fluor 488 NHS ester; Thermo Fisher Scientific for mEGFP). To convert fluorescence intensity to the concentration, FCS was performed in the cells that transiently-express mEGFP alone. Then a calibration curve was obtained by plotting the fluorescence intensity along the concentration. The background fluorescence signal was measured in cells without expressing fluorescent proteins and subtracted.

To measure the concentration of Nups, mEGFP-Nup genome-edited cells in interphase were imaged in 3D using a confocal microscope (LSM780; Carl Zeiss, Oberkochen, Germany) and a 40× 1.2 NA C-Apochromat water immersion objective (Carl Zeiss) at 37 °C in a microscope-body-enclosing incubator, under the following conditions: 21 optical sections, section thickness of 2.0 μm, z-stacks of every 1.0 μm, and the xy pixel size of 0.25 μm. When the NE is not perpendicular to the confocal plane of the 3D stacks, the fluorescence intensity at the NE is nonisotropic in the point-spread function (PSF), which results in underestimation of the signal. To avoid such underestimation, a single plane was selected that contains the largest nuclear area in which the NE is perpendicular to the imaging plane and thus isotropic in the PSF. The fluorescence intensity of Nups was quantified on this single plane using the NE mask with the width of three pixels (0.75 μm) that was generated from a SiR-DNA channel. Background fluorescence intensity was measured in wild-type cells without expressing any fluorescent proteins and subtracted. The Nup fluorescence intensity on the NE was converted to the concentration using the calibration curve generated by FCS above. The number of Nups per square micro-meter was calculated from the concentration and then divided by the NPC density per square micro-meter measured by stimulated emission depletion (STED) microscopy. This absolute quantification of Nup copy number with FCS calibration was done using 47 mEGFP-Nup107 genome-edited cells in interphase. For other mEGFP-Nups genome-edited cells, their Nup fluorescent intensities on the NE were directly compared with the ones of mEGFP-Nup107 genome-edited cells on the same 8-well Lab-Tek Chambered Coverglass, and then their concentrations were determined using the intensity ratios to the mean intensity of mEGFP-Nup107 without using a FCS calibration curve. For Pom121-mCherry, the copy number was quantified independently by performing FCS using Alexa Fluor 568 NHS ester (Thermo Fisher Scientific) to measure the confocal volume and using the cells that transiently-express mCherry alone to convert fluorescence intensity to the concentration.

### Measurement of nuclear pore density by stimulated emission depletion (STED) microscopy

Cells were fixed with 2.4% formaldehyde (Electron Microscopy Sciences, Hatfield, PA) in PBS for 10 min, extracted with 0.4% Triton X-100 (Sigma Aldrich) in PBS for 5 min, and blocked with 5% normal goat serum (Life Technologies, Carlsbad, CA) in PBS for 10 min at room temperature. Subsequently, the cells were incubated overnight at 4°C with a mouse anti-Nup62 (Cat. No. 610497; BD Biosciences, Franklin Lakes, NJ, 1.25 μg/ml) antibody, and then with an Abberior® STAR RED-conjugated anti-mouse IgG (Cat. No. 2-0002-011-2, Abberior GmbH, Göttingen, Germany, 0.5 μg/ml) for 30 min at room temperature. For Extended Data Fig. 3b, anti-GFP (Cat. No. 11814460001; Roche, Basel, Switzerland, 1:200) and anti-Elys (Cat. No. HPA031658; The Human Protein Atlas, 0.5 μg/ml) antibodies, Abberior STAR RED-conjugated anti-rabbit IgG (Cat. No. 2-0012-011-9; Abberior GmbH, 0.5 μg/ml) and Abberior STAR 580-conjugated anti-mouse IgG (Cat. No. 2-0002-005-1; Abberior GmbH, 0.5 μg/ml) were used. After multiple washes in PBS, cells were mounted in Vectashield (Cat. No. H-1500, Vector Laboratories Inc., Burlingame, CA). Super-resolution imaging was performed on a Leica SP8 3X STED microscope as described in a previous report^12^. The images were taken with a final optical pixel size of 20 nm, z-stacks of every 250 nm, and the optical section thickness of 550 nm. Images were filtered with a Gaussian filter (kernel size: 0.5 × 0.5 pixel) for presentation purposes. The shrinkage of the nucleus caused by formaldehyde fixation and/or Vectashield mounting was quantified by comparing the volume of the nuclei of live cells with the ones of fixed cells. The shrinkage was 9.1 ± 2.6% (the average and standard error, N = 36 cells). The NPC density was corrected for the nuclear shrinkage for the calculation of Nup copy number per NPC in Fig. 1d.

To quantify NPC density, the raw STED data were processed in ImageJ^41^ with a mean filter (kernel size: 2 × 2 pixels) and a sliding paraboloid (radius: 5 pixels) for background subtraction. Detection of central peak positions for individual NPCs was carried out with the plugin TrackMate^42^, using DoG detector and adjusting the detection threshold as the spot diameter size. The resulting 3D NPC coordinates were used to visualize and determine flat and curved regions of the nucleus. Using this map, circular and ellipsoidal ROIs could then be selected in the flatter parts containing central NPC positions within the Z-depth of approx. 500 nm, which corresponds to 2–3 microscopic slices in the images. The remaining signal outside the ROIs, as in curved regions or cytoplasmic structures were discarded from further analysis. NPC densities were calculated for each cell separately by dividing the number of NPCs within the selected ROIs by the corresponding ROI areas. For each cell line, the values were combined to calculate the mean and median NPC density values.

### Three-dimensional (3D) STED microscopy for visualizing single nuclear pore assembly intermediates

Cells were coated on 18 × 18 mm #1.5 square coverslips and synchronized by double thymidine arrest. After 10 hours release from the second thymidine treatment, cells were fixed with 2.4% formaldehyde in PBS for 15 min, extracted with 0.25% Triton X-100 (Sigma Aldrich) in PBS for 15 min, and blocked with 2% Bovine Serum Albumin (Cat. No. A2153; Sigma Aldrich) in PBS for 30 min at room temperature. Subsequently, the cells were incubated overnight at 4°C with rabbit anti-Tpr (Cat. No. HPA019661; The Human Protein Atlas, 1:100) and a GFP-nanobody (FluoTag®-X4 anti-GFP nanobody Abberior® Star 635P; Cat. No. N0304-Ab635P-L; NanoTag Biotechnologies, Göttingen, Germany, 1:50), and then with an Alexa Fluor 594 goat anti-rabbit IgG (Cat. No. A-11037; Life Technologies, 1:250) for 30 min at room temperature. After multiple washes in PBS, cells were mounted in ProLong™ Gold Antifade Mountant (Cat. No. P10144; Invitrogen). Super-resolution imaging was performed on an Abberior STED/RESOLFT Expert Line microscope (Abberior GmbH). Samples were imaged with an UPlan-S Apochromat 100× 1.4 NA oil-immersion objective on an IX83 stand (Olympus, Tokyo, Japan). Stimulated depletion was performed using a 775nm pulsed laser (40 MHz) in combination with 594 and 640 nm pulsed excitation lasers in line switching mode. Fluorescence signal was detected using two separate Avalanche photo diodes with bandpass filters of 605–625 and 650–720 nm. The images were taken with a final optical pixel size of 35 nm, z-stacks of every 200 nm, and the optical section thickness of 1000 nm. For presentation purposes, images were filtered with PureDenoise plugin^43^ in ImageJ, and lines with the width of 175 nm were drawn along the nuclear envelope, flattened and shown in the figures.

### Quantification of Nup copy number in the cytoplasm and the nucleoplasm as well as in non-core and core regions of the NE

Mitotic cells were imaged and monitored from anaphase onset for two hours in 3D by confocal microscopy. The microscopy setup and the imaging conditions are described above. Time-lapse imaging for mEGFP-tagged Nups was performed every 30 sec. Photobleaching was negligible and thus not corrected. Time-lapse imaging for Pom121-mCherry was carried out every 60 sec, and photobleaching was corrected by measuring a fluorescence signal decay in a neighboring cell in the same field of view. Visualization of the chromosome surface in 3D was done in the Amira software package^44^.

To measure the Nup accumulation on the NE, single planes were selected that contain the largest nuclear area at individual time points to avoid underestimation of the signal as mentioned earlier. The Nup intensity was quantified on the NE mask with the width of 0.75 μm that was generated from a SiR-DNA signal at each time point. Except for Nup107 and Seh1, the Nup signal in the cytoplasm and nucleoplasm was measured and used as background. For Nup107 and Seh1, only the cytoplasmic signal was used as background because of their localization at kinetochores. These background values were quantified at individual time points and subtracted from the Nup intensities on the NE. The measured Nup intensity was converted into the concentration and then multiplied with nuclear surface area to calculate the total number of the Nups on the NE. For Nup153 and Pom121, we did not convert the fluorescence intensity to the concentration as the cell lines were not fully validate to be homozygously-tagged.

The Nup copy number was also calculated in the cytoplasm and the nucleoplasm during first two hours after anaphase onset. The cytoplasm mask was created by subtracting a mask of the nucleus generated from a SiR-DNA signal from a mask of the whole cell generated from a mEGFP-Nup signal. The mask for the nucleoplasm was created by eroding three pixels of the nuclear mask generated from a SiR-DNA signal. The Nup fluorescence intensity was quantified on these cytoplasmic and nucleoplasmic masks over time and then converted into the concentration. To calculate the total number of Nups in the cytoplasm and the nucleoplasm, the measured concentration was multiplied by the volume of the respective compartments. For the cytosol volume at metaphase, we have quantified it using the cytosolic signal of the Nups. The cytosolic GFP signal of some of the Nups (e.g. Nup205, Nup214, Tpr, Nup358) were too low to precisely segment the cytosol in the z-slices close to the glass surface due to the relatively high background fluorescence signal. Therefore, we measured the cell volume using the brightest Nup62-mEGFP cell line (the volume was 4900 μm^3^ with the s.d. of 330 μm^3^, from 8 cells), and for the other Nup cell lines, we measured the cytosol area on a middle z-slice plane and calculated the ratios to the one of the Nup62 cell line (the ratios were 0.97, 0.97, 1.09, 1.14, 1.03, 1.07, and 1.02 for Nup107, Seh1, Nup205, Nup93, Nup214, and Nup358, respectively). From the measured volume of the Nup62 cell and the ratios of cytosol area to the Nup62 cell, we have estimated the volume of other cell lines (the volume was 4670, 4660, 5550, 6030, 5110, 5420, and 5080 μm^3^ for Nup107, Seh1, Nup205, Nup93, Nup214, and Nup358, respectively). For the cytoplasm volume after anaphase, we utilized the data that was measured previously in histone H2b-mCherry expressing HeLa cells using fluorescently-labelled dextran^17^. Assuming that the cell volume changes in the same degree during mitosis exit as the H2b-mCherry cell, we calculated the cytoplasmic volume of the Nup cell lines using the ratio to the volume at metaphase (5300 μm^3^ for the H2b-mCherry cell^17^). The nucleoplasmic volume was quantified in each mEGFP-Nup knock-in cell line using a SiR-DNA signal at every time points as described previously^12^.

Core regions were predicted on the NE based on a previously described protocol using the core marker Lap-2α^12^. Briefly, nuclear volume was segmented using SiR-DNA fluorescence signals that were processed with a 3D Gaussian filter and a multi-level thresholding. Nuclear volume was then divided into inner and outer volumes using the cutting plane that was constructed from the largest eigenvector and the second one orthogonal to the first vector of the pixel coordinates of the nuclear volume. Surface area of each nucleus was calculated and utilized to adjust the size of the inner and outer-core regions at individual time points. The previously defined criteria for being core and non-core regions^12^ was applied. The position of inner and outer core was determined with respect to the intersection point of the largest eigenvector on the nuclear surface. The core region prediction was done in MATLAB (The MathWorks, Inc.).

### Mathematical modeling for the nuclear pore assembly kinetics

Previous EM data showed that within 2 hours after the onset of anaphase, postmitotic assembly is the dominant process in the non-core region, whereas slower interphase assembly predominates in the core region^12^. Assuming that this relation is also reflected in the live-cell Nup dynamics, we derived a mathematical model. We assumed that the observed total fluorescence intensity in the non-core, *n*(*t*), and core region, *c*(*t*), at time-point *t* after anaphase onset is a linear combination of the postmitotic, *pm*(*t*), and interphase assembly, *ip*(*t*), according to

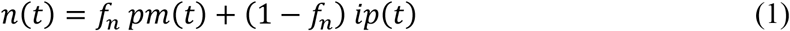

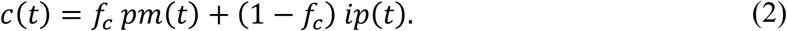

The fraction of postmitotic assembly in the non-core and core regions are denoted *f_n_* and *f_c_*, respectively. To test this assumption and obtain an estimate of the fractions, we used a model that accounts for the observed sigmoid-like kinetics

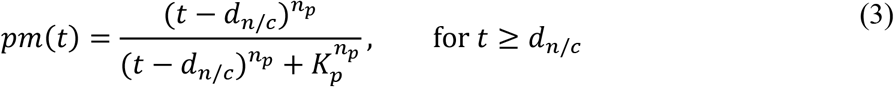

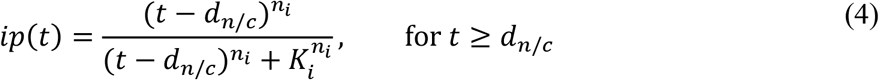

and *pm*(*t* < *d_n/c_*) = *ip*(*t* < *d_n/c_*) = 0. The parameters *n_p_, K_p_* and *n_i_*, *K_i_*, characterize the postmitotic and interphase kinetics, respectively. The parameter *d_n/c_* accounts for an additional delay in NPC initiation. In the non-core region we assumed *d_n_* = 0, in the core region due to the presence of kinetochore microtubule fibers^28^, *d_c_* > 0. The model is used to derive the underlying post-mitotic and interphase assembly kinetics. Using Eqs. **Error! Reference source not found.–Error! Reference source not found**. we obtain

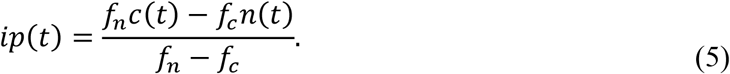

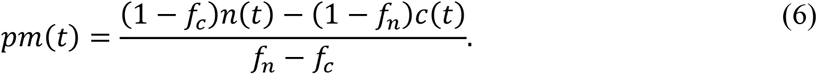

#### Parameter estimation

For each Nup, there are two parameters that define the interphase (*n_i_*, *K_i_*) and postomitic (*n_p_*, *K_p_*) assembly, respectively. The fractions *f_n_* and *f_c_* and the delay *d_c_* in the core region were estimated globally for all Nups. In total, we have 43 parameter, 4*10 = 40 parameters describing the assembly kinetics and 3 global parameters, fitted to 4446 data points.

In detail, to find the model parameters we minimized the mean squared distance between data and model for all the time points *M*

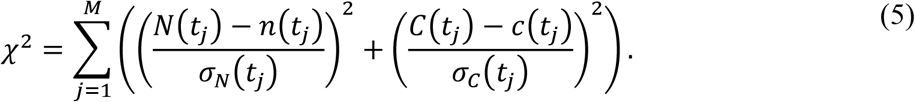

*N*(*t_j_*) and *C*(*t_j_*) are the mean background-subtracted and normalized fluorescence intensities in the non-core and core region with standard deviation *σ_N_*(*t_j_*) and *σ_C_*(*t_j_*), respectively, at time point *t* = *t_j_*. We subtracted a background computed from the average of the first 3 time points. The data were normalized with the average value between 100 and 120 min after anaphase. In a first step, for each protein/cell-line, we estimated the postmitotic fractions in the core and non-core region and the kinetic parameters. Overall, we computed 61 parameters (6 parameters per protein plus one delay parameter). For the postmitotic fractions, we obtained on average *f_n_* = 0.857 [0.76, 0.95] and *f_c_* = 0.295 [0.17, 0.4], where the number in brackets indicates the 95% confidence interval estimated using the profile likelihood method^45^. Importantly, the obtained postmitotic fractions are well in agreement with the previously reported estimates obtained from EM-data^12^. The delay in pore formation between core and non-core region was estimated by systematically varying *d_c_* from 0 to 6 min in steps of 1 minute. A value of *d_c_* = 2 min, gave optimal result. In a second step, we used the previously estimated average postmitotic fractions, *f_n_* and *f_c_*, and *d_c_* and recomputed the kinetics parameters for each protein. The model with reduced parameters well agrees with the data (Extended Data Figs. 4 and 5, R^2^ > 0.99). To verify if the choice of common postmitotic fractions for all Nups is valid, we computed the Baysian information criterion (BIC)^46^. The difference in BIC between the model with reduced parameters, 43 parameters, compared to the full model, 61 parameters, was −7, indicating that the model with reduced parameters is justified. The obtained parameter values are listed in Supplementary Table 2.

#### Median assembly time and duration of assembly

One can think of an assembly curve as a cumulative distribution function and its derivative as the corresponding probability density function (PDF) for binding events. The median assembly time, i.e. the time where 50% of binding events have occurred, is *K_i_* and *K_p_*, for the interphase and postmitotic assembly respectively. We consider the time intrinsic to the assembly mechanism and independent of the initiation delay *d_c_*. We further define the duration of assembly as the time interval where 80% of binding events occur, i.e. from a fraction of *α*_1_ = 0.1 up to *α*_2_ = 0.9, one obtains (see also Extended Data Fig. 6a)

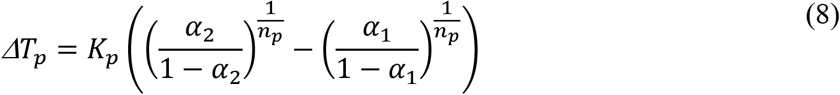

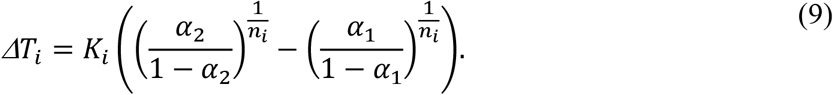

The duration of assembly quantifies the width of the binding events PDF.

In Extended Data Fig. 6c we see a strong positive correlation between median assembly time and duration of assembly for the postmitotic assembly. This suggests a sequential assembly mechanism. The rationale is that for a strict sequential pathway, the binding events PDF for subsequent Nup is a convolution of all previous binding events PDF and so will broaden for later binding proteins. For example, if an early protein has an assembly duration of 1 hour, within this time window binding sites for the subsequent protein will continue to appear. Therefore, the subsequent protein in the sequence will also accumulate for at least 1 hour.

For the simplified case of irreversible sequential assembly and linear rate constants, a positive correlation can be demonstrated. This correlation is independent on whether the initiation is synchronous or spread within a time-period. For simplicity reasons, we omit the fact that each nucleoporin binds in multiple copies to the NPC. A nucleoporin P*_i_* binds with rate constant *κ_i_* to a NPC intermediate X_*i*-1_ according to following reaction scheme

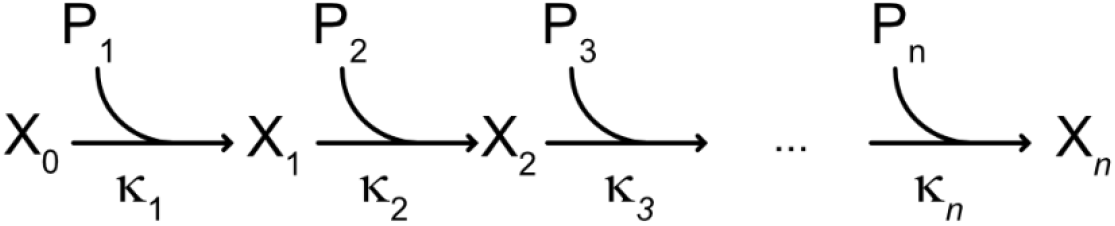

The corresponding system of ordinary differential equations is

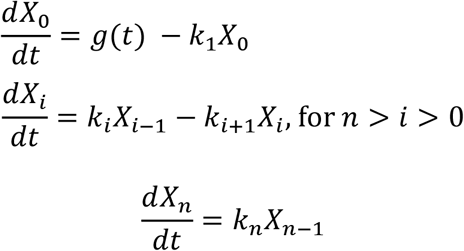

with *X_i_*(0) = 0, 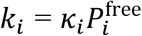, and an excess of free nucleoporin 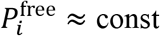. The function *g*, accounts for the appearance of NPC initiation sites after anaphase. We assume that NPC initiation is completed within a finite time after anaphase and neglect the slowly and continuous appearance of pores in G1, i.e. 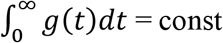. The amount of *i*th Nup bound to a complex is 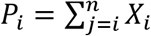 and its time derivative 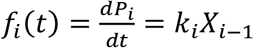 is proportional to the binding events PDF. We can quantify the assembly time *τ_i_* and assembly duration *θ_i_* by the mean and standard deviation of the binding events PDF, respectively^47^,

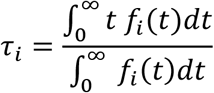

and

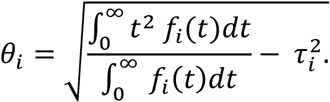

After integration by parts one obtains

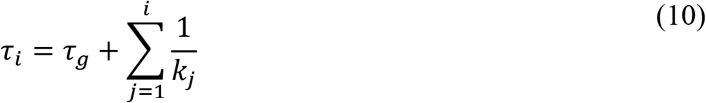

and

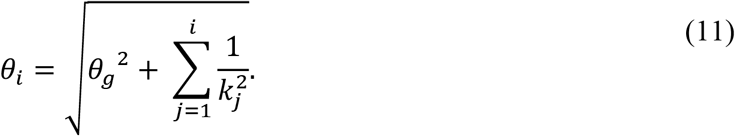

Where 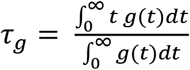 and 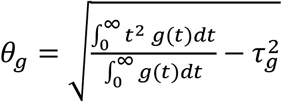 is the mean and standard deviation of the initiation function, respectively. From Eqs. 10–11, it is clear that *τ*_*i*+1_ > *τ_i_* and *θ*_*i*+1_ > *θ_i_* for any parameter combinations and independently on how synchronous the initiation of pores is. This shows that for a strictly sequential pathway a linear correlation between assembly time and duration is expected.

### Integrative modeling of the NPC assembly pathway

A model of the assembly pathway is defined by a series of static structures, including a static structure at each sampled time point along the assembly process. Therefore, we model the NPC assembly by first modeling static structures at each time point, independently from each other. We then enumerate alternative assembly pathways and rank them based on the static structure scores and plausibility of transitions between successive static structures.

#### Integrative modeling of static structures at each time point

The static structures are modeled by standard integrative structure modeling^32^, as follows.

#### Representing a static structure model

The time points correspond to times with available ET protein densities^11^: 5min, 6 min, 8 min, 10 mins, and 15 min after anaphase onset. We divide the mature NPC structure (PDB 5a9q, 5ijo) into eight spokes and further divide each spoke into a set of rigid subcomplexes, including the Y-complex, the inner ring Nup205-Nup155-Nup93 subcomplex, the inner ring Nup93-Nup188-Nup155 subcomplex, and the central channel Nup62-Nup58-Nup54 subcomplex. For each domain, we coarse-grained the structure by grouping 10 consecutive amino acid residues into a single bead at the center of mass of those residues. Each subcomplex is represented as a rigid body. The NE is represented as a fixed toroid surface embedded in two parallel planes. Thus, the variables of the model include the Euclidean coordinates of the Nup subcomplexes and the copy number of each Nup subcomplex.

We set the inner pore diameter and minor radius of the pore at each time point to the mean of previously determined NE cross sections^11^ with a pore diameter of 51.5 nm, 58.4 nm, 72.7 nm, 84.6 nm, 79.8 nm, and 87 nm; and minor radius of 21.4 nm, 21.2 nm, 21.5, 20.3 nm, 17.1 nm and 15 nm for time points at 5min, 6min, 8min, 10min, 15min, and the mature pore, respectively.

#### Scoring a static structure model

The copy numbers of the NPC subcomplexes at each time point were restrained by a Gaussian function with mean and variance determined by the single-cell traces presented in this study. The relative likelihood of a set of copy numbers is proportional to the product of individual Gaussian likelihoods.

Distances between pairs of Nups that are in contact with each other in the native NPC structure^29,30^ were restrained by a harmonic Gō-like model^48^. Inter-subcomplex contacts within 5 nm in the mature structure were restrained by a harmonic function (strength 0.01 kcal/mol Å). Each Gō-like scoring term was scaled at each time point, from zero at the first time point to full strength at the mature pore time point. Distances between all pairs of Nups were also restrained by a harmonic excluded volume restraint (strength 0.01 kcal/mol Å). Proximity between Nup domains containing a membrane interacting ALPS-motif and the NE was restrained by a harmonic term (strength 0.1 kcal/mol Å), based on their sequences. Overlap between the Nups and NE surface was avoided by imposing a harmonic repulsion between the Nups and NE surface (strength of 0.01 kcal/mol Å).

The shape of a static structure was restrained by a correlation coefficient between the model and ET protein density^11^. The forward model density was represented by fitting each Nup subcomplex with a Gaussian mixture model of two components per subcomplex copy using the *gmconvert* utility^49^. Similarly, the ET protein densities at each time point were represented with a Gaussian mixture model with 150 components fit to the experimental density.

#### Sampling static structure models

A state of the NPC at any given time point is defined by the copy numbers and coordinates of its components. Only copy number assignments and structures consistent with the C-8 symmetries were sampled. The assumption that the subcomplexes preform with 8-fold multiplicity before assembling into the NPC is supported by our previous electron microscopy analysis^11^. The averaged electron tomograms of the intermediates demonstrated 8-fold symmetry of the outer-ring complex at 5, 6, 8, 10, and 15 min after anaphase onset and 8-fold symmetry of the inner-ring complex at 10 and 15 min after anaphase onset^11^, indicating that the majority of intermediates have approximate 8-fold symmetry for the outer- and inner rings. However, there are likely to be variations in the copy numbers at the single pore level. We only sampled structures for the top 20-scoring Nup copy number combinations. Each sampling started with the mature pore structure, followed by applying 10^6^ Monte Carlo moves. These moves included rotational and translational perturbations to each Nup subcomplex, drawn from a uniform distribution in the range from −0.04 to +0.04 radians and from −4 to +4 Å, respectively.

#### Modeling the assembly pathway

With the static structure models in hand, we connect them into complete alternative assembly pathways, as follows.

Each pathway is represented by a static structure at each sampled time point, starting with t = 5 min and culminating in the native structure; we do not model the completely disassembled NPC. The score of a pathway is the sum of the scores for the static structures on the pathway (defined above) and transitions between them. A transition score is uniform for all allowed transitions. A transition between two successive static structures is allowed if the subcomplexes in the first structure are included in the second structure. All possible pathways were enumerated, scored, and ranked. The best-scoring pathways were extracted for further analysis (Fig. 5). Molecular visualization was performed with UCSF ChimeraX^50^.

### Immunofluorescence for visualizing Nup155 during assembly

Cells were treated, fixed and blocked as described in the previous section (STED microscopy for visualizing single nuclear pore assembly intermediates). Afterwards, cells were incubated overnight at 4°C with rabbit anti-Nup155 (Cat. No. HPA037775; The Human Protein Atlas, 1:100), and then with an Abberior STAR635P Goat anti-rabbit IgG (Cat. No. ST635P-1002-500UG, Abberior GmbH, 1:250) for 30 min at room temperature. After multiple washes in PBS, cells were mounted in ProLong™ Gold Antifade Mountant with DAPI (Cat. No. P36941, Invitrogen). Imaging was performed in 3D using a confocal microscope (LSM780; Carl Zeiss) and a 63× 1.4 NA Plan-Apochromat objective (Carl Zeiss) under the following conditions: 31 optical sections, section thickness of 0.7 μm, z-stacks of every 0.39 μm, and the xy pixel size of 0.13 μm.

### Sample size determination and statistical analysis

For quantitative imaging in Fig. 1a, d, the data were from 4, 4, 4, 2, 3, 2, 3, and 2 independent experiments for Nup107, Seh1, Nup205, Nup93, Nup62, Nup214, Tpr and Nup358, respectively. STED imaging in Fig. 1b, c and Fig. 4 were from 1 and 2 independent experiments, respectively. For dynamic quantitative imaging in Fig. 3, the data were from 4, 4, 4, 2, 3, 4, 3, 2, 2, and 4 independent experiments for Nup107, Seh1, Nup205, Nup93, Nup62, Nup214, Tpr, Nup358, Nup153, and Pom121, respectively. Southern blotting in Extended Data Fig. 1 and immuno-fluorescence microscopy in Extended Data Fig. 8 are from single experiments. Statistical analyses were performed only after all the data were taken. Sample sizes for each experiment are indicated in figure legends. Sample sizes were based on pilot experiments to determine the number of cells required to observe stable population averages with high Pearson’s correlation between replicates. Videos of dividing cells with rotating nuclei are removed from the analysis, because we cannot properly assign the non-core and core regions.

### Data availability

Numerical and statistical source data for Figs. 1c, d, 2b, 3a, b, 4c, and Extended Data Figs. 3a, 4b– d, 5, 9e are provided online. Fluorescence images were deposited to the Image Data Resource (IDR; https://idr.openmicroscopy.org/) under accession number idr0115. Our integrative spatiotemporal model of the postmitotic assembly of the human NPC is deposited to the nascent database of integrative structures PDB-Dev (https://pdb-dev.wwpdb.org/) under accession number PDBDEV_00000142. The full scan of the blot shown in Extended Data Fig. 9b is provided online.

### Code availability

All source code is accessible on Github repository (for quantifying Nup intensity in core and non-core regions: https://github.com/mjh1m22/Quantitative_map_nups_Otsuka_Nature_2022; for decomposing postmitotic and interphase assembly kinetics; https://github.com/manerotoni/npc_assembly_Otsuka_2022). *Integrative Modeling Platform* (IMP) is an open-source program freely available under the LGPL license at http://integrativemodeling.org; all input files, scripts, and output files are available at http://integrativemodeling.org/npcassembly.

## Acknowledgments

We thank the EMBL Advanced Light Microscopy Facility (ALMF) for their support in STED super-resolution microscopy; Sebastian Schnorrenberg for help with 3D-STED; Andreas Brunner for help with automated live-cell imaging; Ben Webb for help with IMP and Josh Baker-Lepain for managing the Wynton computer cluster at QBI@UCSF. This work was supported by grants from the Baden Wuerttemberg Foundation (J.E. and A.S.), NIH/NIGMS R01GM083960 (A.S.), NIH/NIGMS P41GM109824 (A.S.), NIH/NIGMS R01GM112108 (A.S.) and the European Molecular Biology Laboratory (EMBL; S.O., A.Z.P., A.R., M.J.H., M.K., A.C., B.K., J.E.). S.O. and A.R. were further supported by the EMBL Interdisciplinary Postdoc Programme (EIPOD) under Marie Curie Actions COFUND. S.O. was additionally supported by a JSPS fellowship (The Japan Society for the Promotion of Science, postdoctoral fellowship for research abroad).

## Author contributions

SO, JOBT, AS, and JE designed the project. SO performed all the fluorescence microscopy experiments and analyses, except for the experiments in Fig. 4 and Extended Data Figs. 8 and 9. JOBT performed integrative structural modeling. WZ performed 3D-STED, immuno-fluorescence time course microscopy, and live imaging shown in Fig. 4 and Extended Data Figs. 8 and 9. AZP carried out mathematical modeling for the nuclear pore assembly kinetics. AR developed an analysis pipeline for nuclear pore density measurement. MJH established a computational image analysis pipeline to quantify fluorescence intensities in non-core and core regions in 3D time-lapse images. WZ, MK, AC, BK and NRM generated and validated genome-edited cell lines. AS and JE supervised the work. SO, JOBT, AS, and JE wrote the paper. All authors contributed to the analysis and interpretation of data and provided input on the manuscript.

## Additional information

The manuscript contains supplementary material (Extended Data Figs. 1 to 9, Extended Data Table 1, Supplementary Figs. 1 and 2, and Supplementary Tables 1 and 2). Correspondence and requests for materials should be addressed to S.O. or J.E.

