## Extended Data Figs. 1 to 9; Extended Data Table 1 for "A quantitative map of nuclear pore assembly reveals two distinct mechanisms"

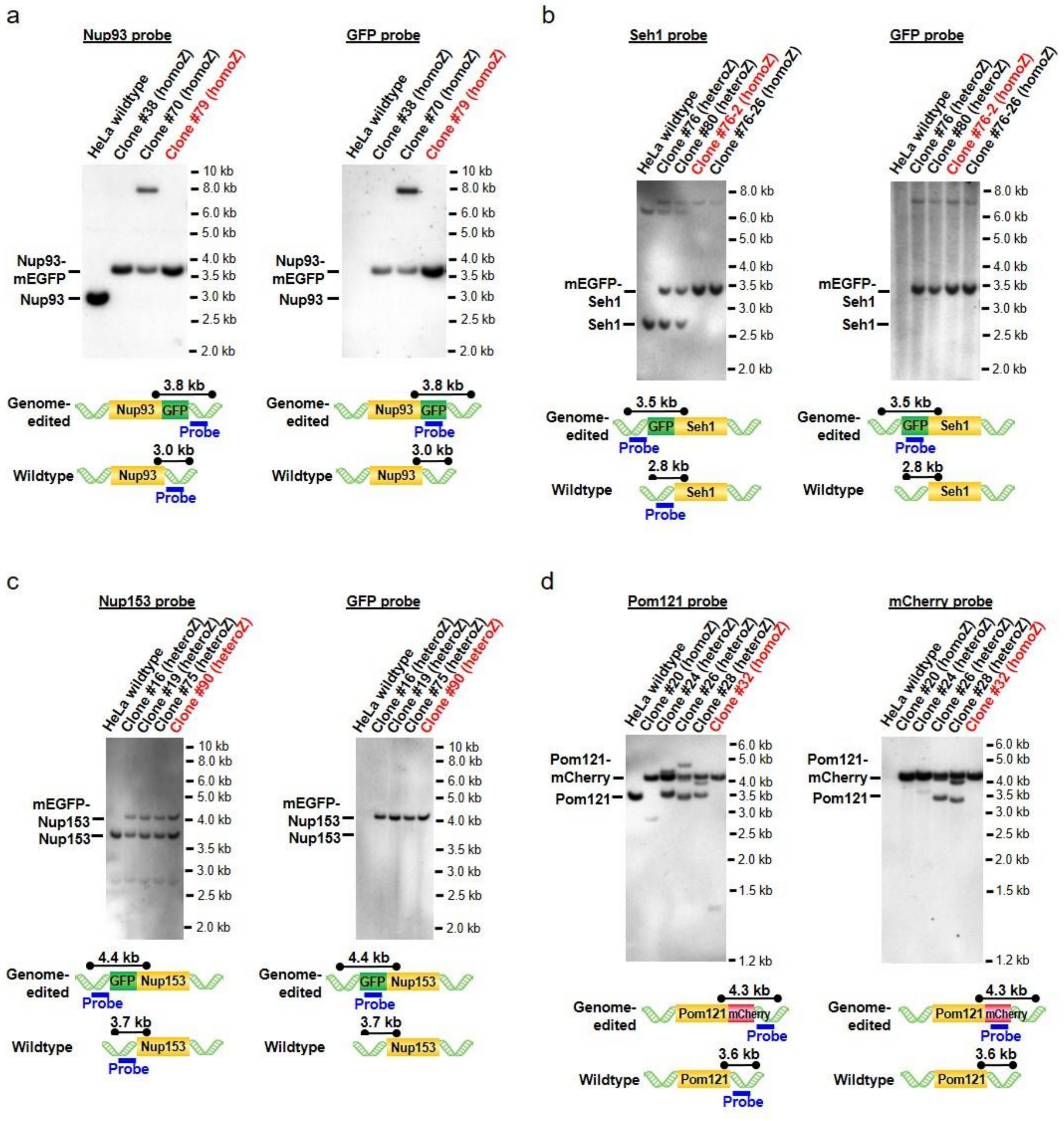


**Extended Data Fig. 1 | Southern blotting of genome-edited cell clones.** Genomic DNA of each cell clone was digested with restriction enzymes and the fragments were detected by probes for Nup93 (**a**), Seh1 (**b**), Nup153 (**c**), and Pom121 (**d**), as well as GFP (**a**–**c**) and mCherry (**d**). The size of the fragments and the probe-binding regions are illustrated in the bottom panels. The clones indicated in red were used for this study. We regarded the clone #32 of Pom121-mCherry as subhomozygous because the protein abundance was much less than expected although the blot indicated homozygous tagging. The following Nup cell lines were validated to be homozygously-tagged in previous reports (Tpr, Nup214 and Nup358[17]; Nup107[12]; Nup205[11]; Nup62[18]).

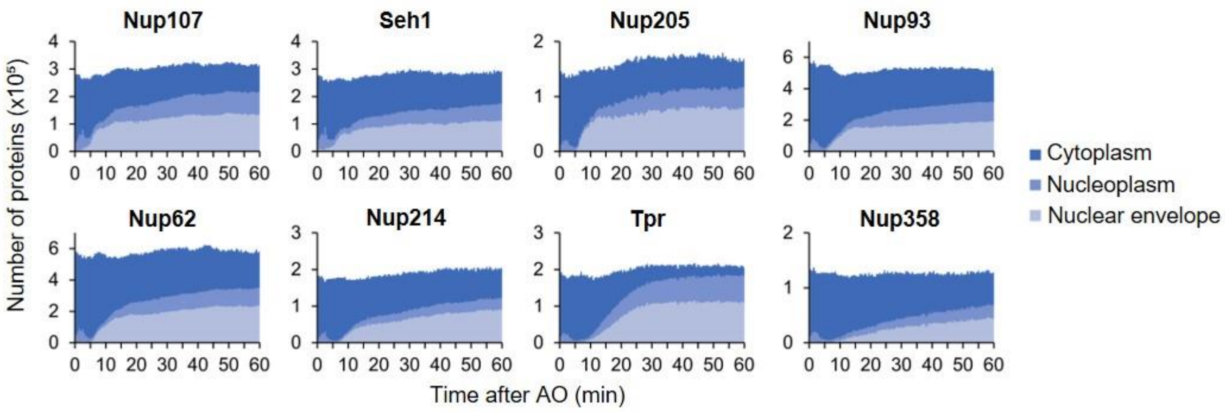


**Extended Data Fig. 2 | NPC assembly relies on and consumes almost half of the material inherited from the mother cell within one hour after mitosis.** FCS-calibrated 3D confocal microscopy was performed as in Fig. 2, and the Nup copy number was quantified in whole dividing cells (for details see Methods). The number of Nups in cytoplasm (dark blue), nucleoplasm (medium dark blue) and nuclear envelope (light blue) are plotted against time after anaphase onset. The plot is the mean of 15, 20, 13, 14, 22, 22, 19, and 24 cells for Nup107, Seh1, Nup205, Nup93, Nup62, Nup214, Tpr and Nup358, respectively.

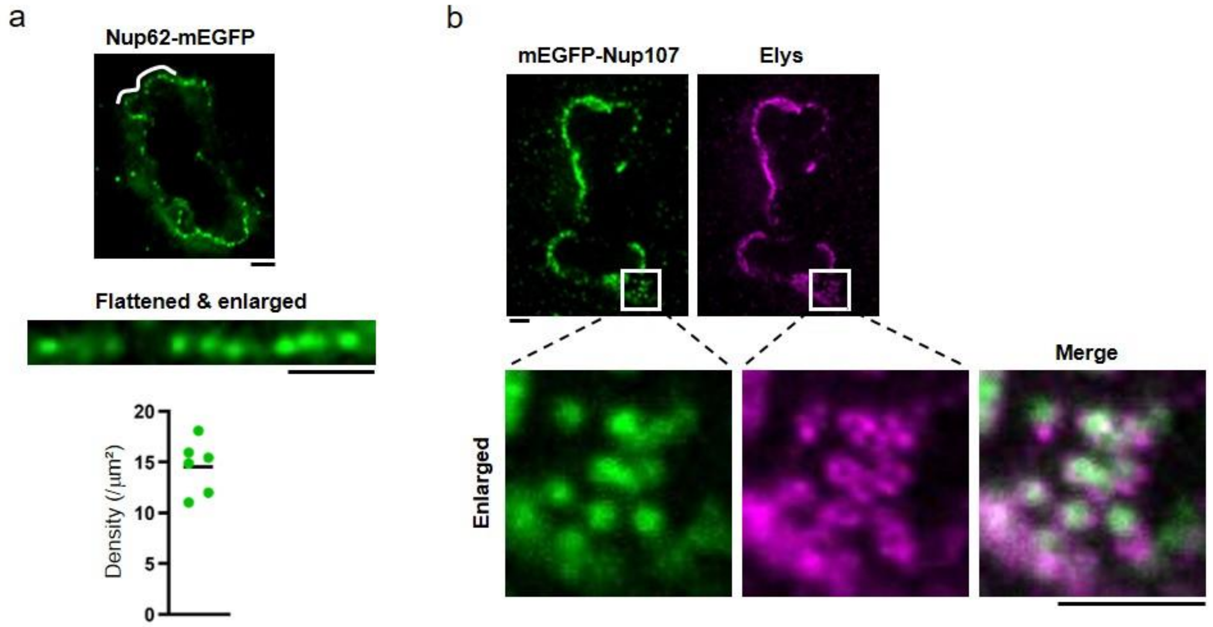


**Extended Data Fig. 3 | GFP-tagged Nups are recruited to the NPCs rather than nonspecifically accumulated on the NE.**

**a,** Three-dimensional stimulated emission depletion (3D-STED) imaging of a Nup62-mEGFP genome-edited cell at early telophase. The region of the nuclear envelope indicated by a white line in the top image is flattened and shown in the bottom. Scale bars, 1 µm. The density of the Nup62 spots was 14.6 ± 2.6 NPCs/$\mu m^2$ (from 6 cells), which is consistent with our previous EM observation of the NPC density of 12–16 NPCs/$\mu m^2$ in early telophase cells[11]. **b,** 2D-STED imaging of a mEGFP-Nup107 genome-edited cell at early telophase that were stained with anti-GFP and anti-Elys antibodies. Enlarged images indicated by white boxes in the top images are shown in the bottom panel. Scale bars, 1 µm.

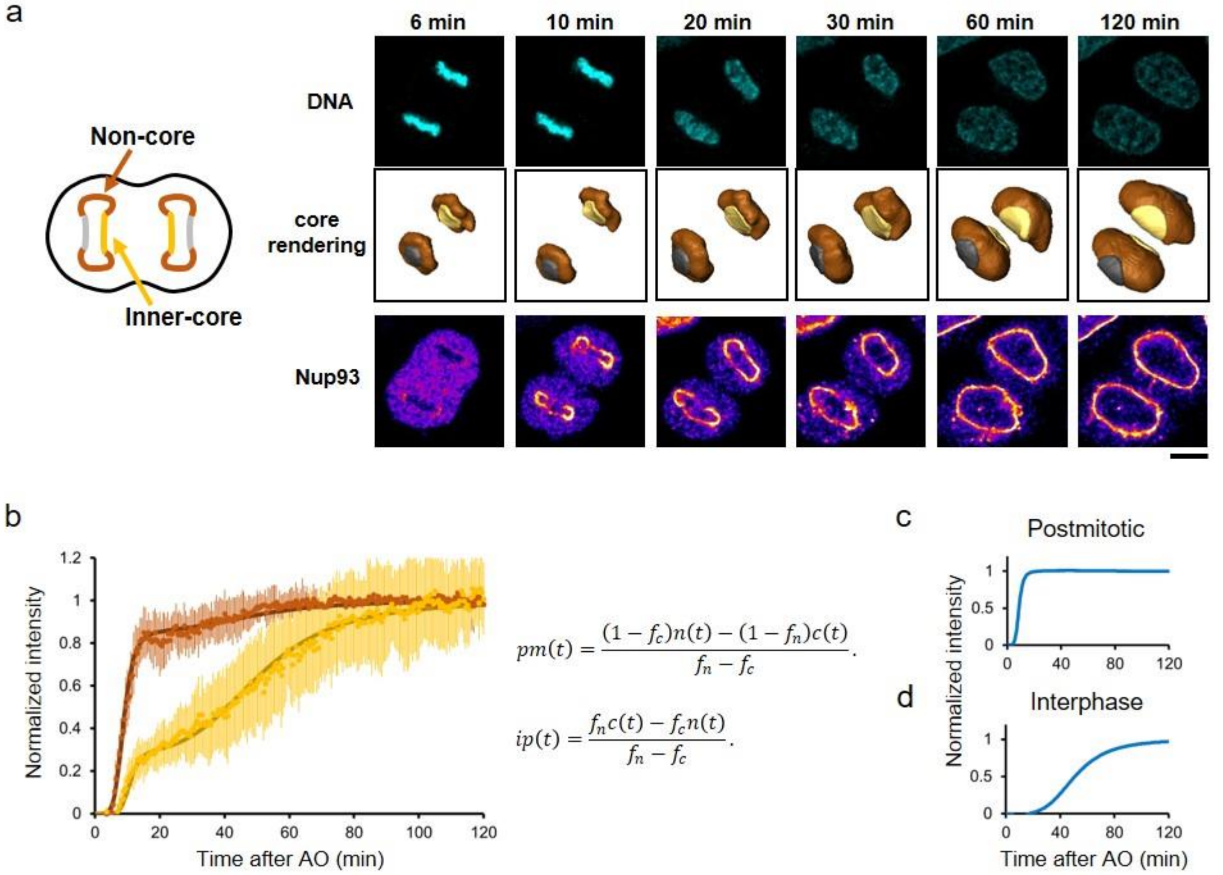


**Extended Data Fig. 4 | Postmitotic and interphase assembly are spatially distinguished for the first hour after mitotic exit and thus their kinetics can be decomposed.**
**a**, Time-lapse 3D imaging of Nup93-mEGFP genome-edited cell. Single confocal sections of SiR-DNA and GFP channels are shown. Images were filtered with a median filter (kernel size: 0.25 × 0.25 µm). Scale bar, 10 µm. Time after AO is indicated. **b**, The fluorescence intensities at non-core (brown) and inner-core (yellow) regions were quantified. Dots represent the average and s.d. of measurements from 14 cells. The intensities were fitted with a sequential model of NPC assembly (bold lines) that allows for different population and rate constants for postmitotic and interphase assembly (described in detail in Methods and Supplementary Table 2). **c**, **d**, Decomposed kinetics of postmitotic (**c**) and interphase (**d**) assembly from (**b**).

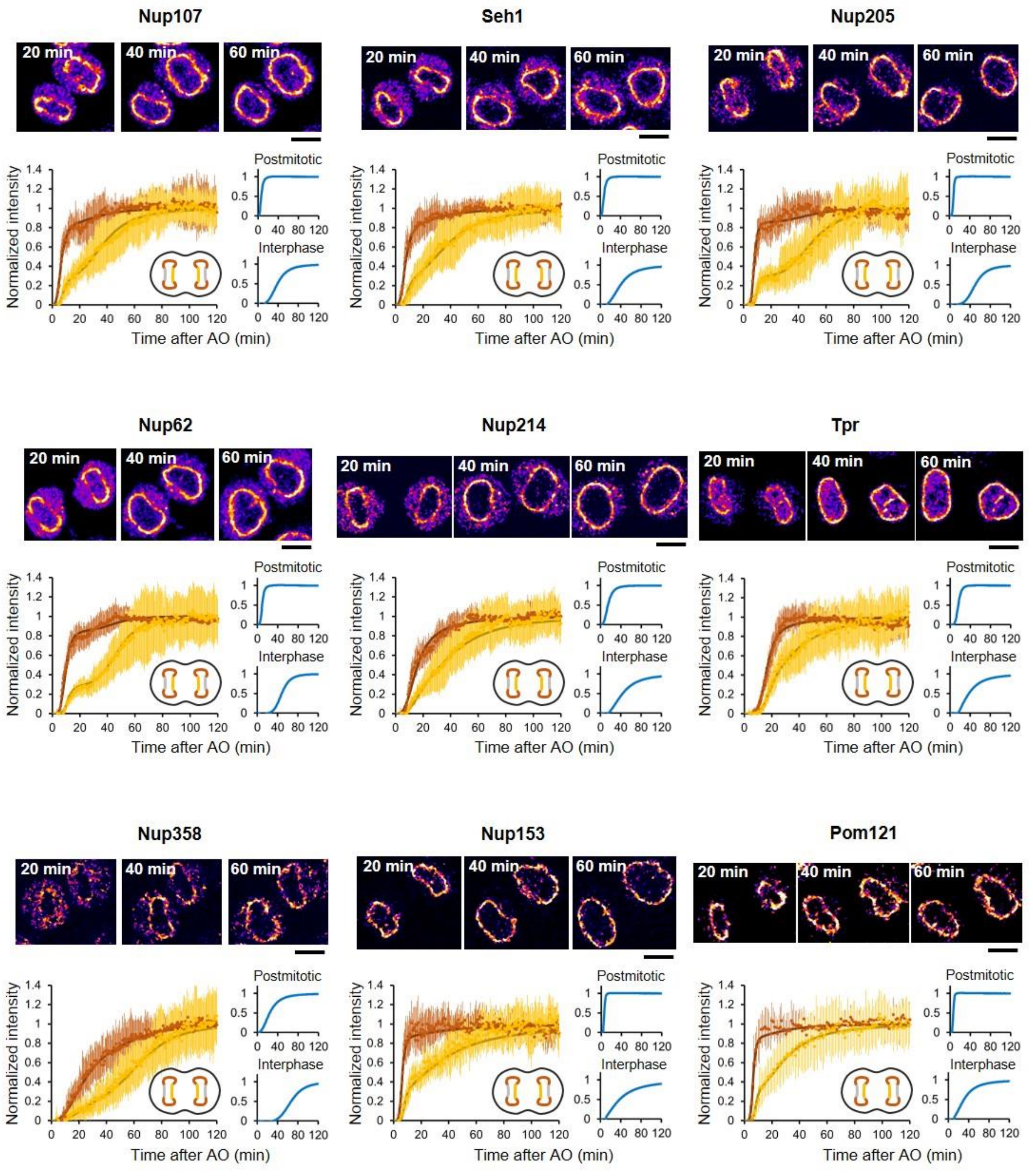


**Extended Data Fig. 5 | Kinetic decomposition of the two assembly processes for each nucleoporin.**

The fluorescence intensities at non-core (brown) and inner-core (yellow) regions were plotted and fitted with a sequential model of NPC assembly (bold lines) as in Extended Data Fig. 4. Dots represent the average and s.d. of measurements from 15, 20, 13, 14, 22, 22, 19, 24, 14 and 13 cells for Nup107, Seh1, Nup205, Nup93, Nup62, Nup214, Tpr, Nup358, Nup153, and Pom121, respectively. Single confocal slices of cells at 20 min, 40 min, and 60 min after AO are shown. Images were filtered with a median filter (kernel size: 0.25 × 0.25 μm) for presentation purposes. Scale bars, 10 μm.

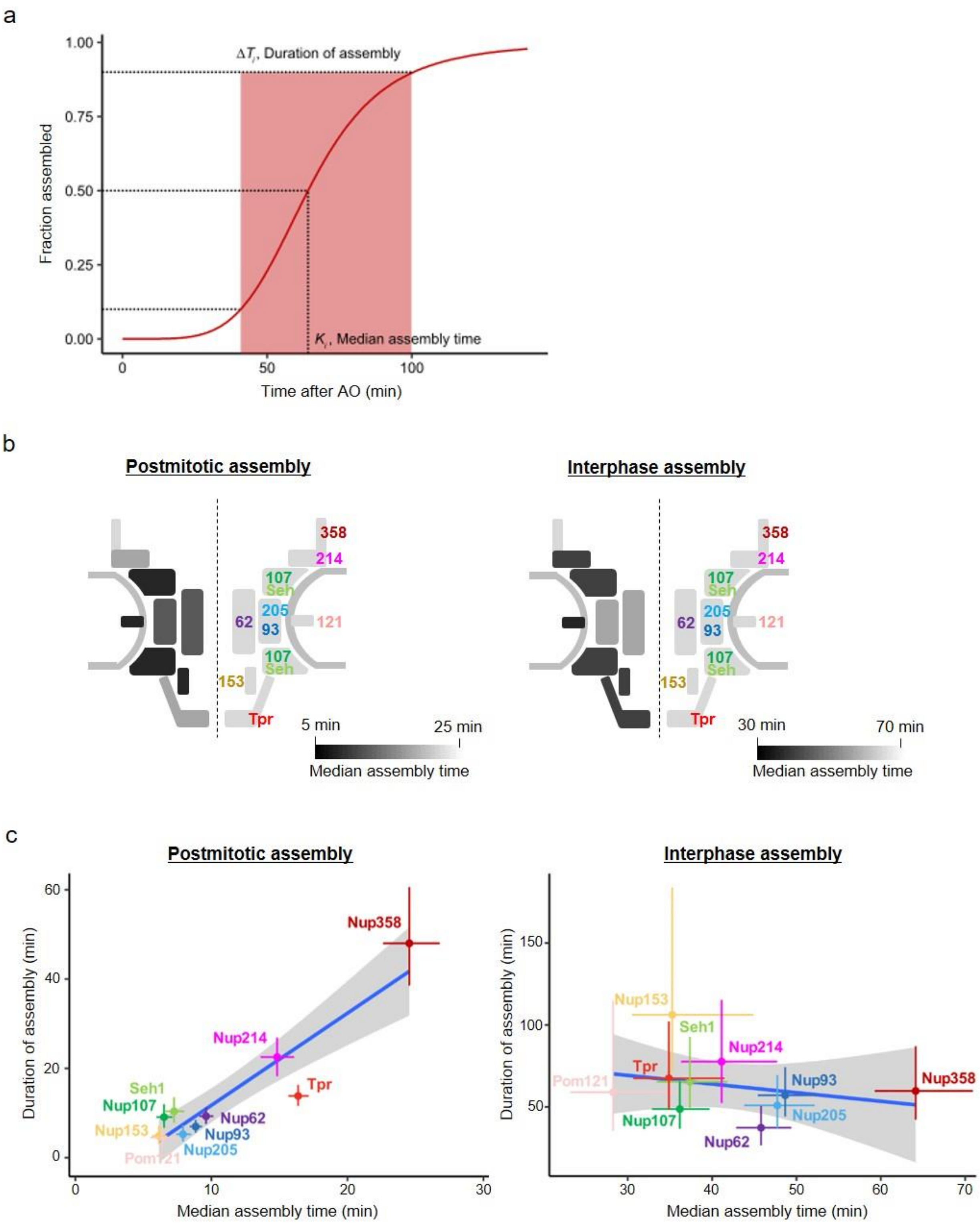


**Extended Data Fig. 6 | Computed parameters for the nucleoporin assembly kinetics and the remarkable difference between postmitotic and interphase assembly.**
**a**, An example of computed quantities (Nup358, interphase assembly). **b**, Illustrations of median assembly time of nucleoporins in postmitotic (left) and interphase (right) assembly superimposed onto representing NPC structures. The median assembly time of individual nucleoporins are displayed on a pseudo-colour scale. **c**, Plots of Nup assembly duration along the median assembly time for postmitotic and interphase assembly pathways. The crosses indicate the 95%

confidence intervals. The dots are the mean values. Values are listed in Supplementary Table 2. The long straight line shows the result of a linear regression to the mean values. The gray area is the 95% confidence interval of the linear regression. Theoretically, for a sequential assembly mechanism where late steps depend on early steps, the observed ensemble kinetics of a late binding protein must contain the history of all previous events (see Methods for details). Indeed, postmitotic assembly showed a strong positive linear correlation, indicating a sequential assembly mechanism, which for example implies that Nup62 is incorporated into the NPC before Tpr can bind.

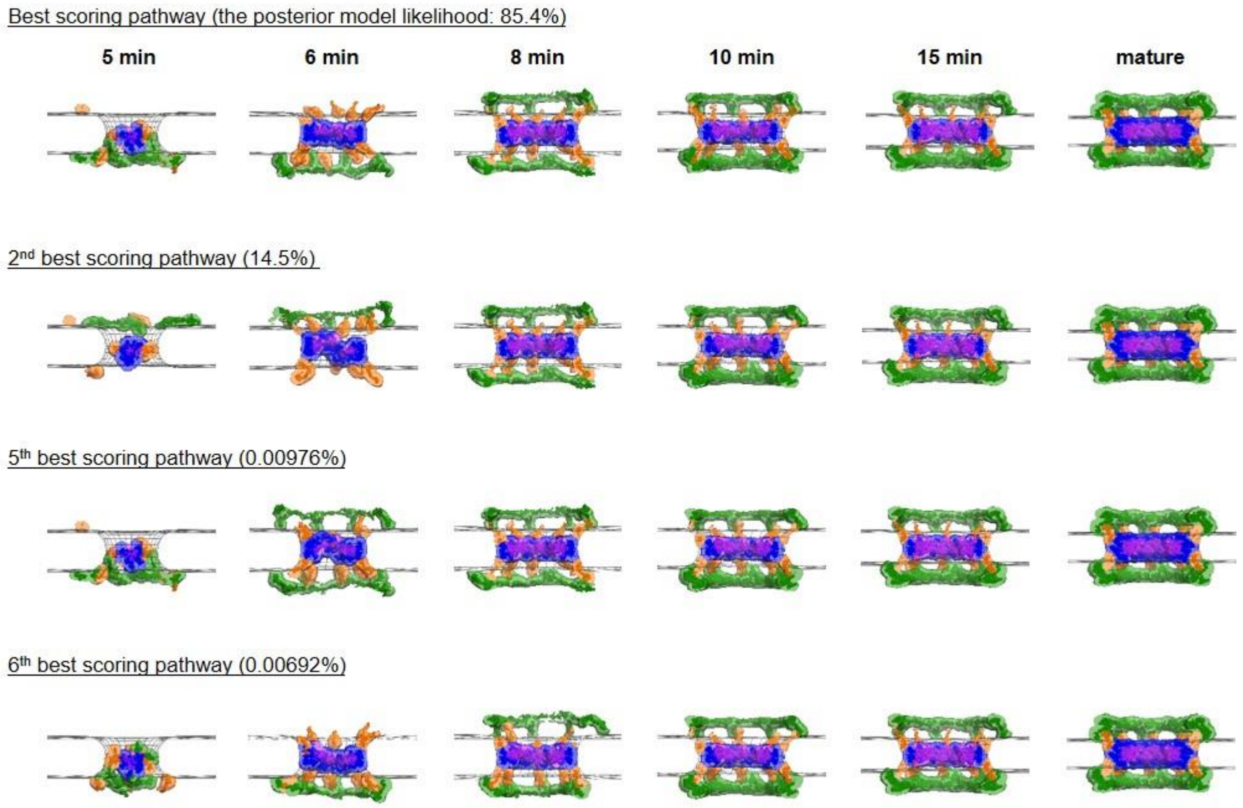


**Extended Data Fig. 7 | The variations of the integrative models for postmitotic assembly pathway.**

Detailed views of the best and lower-scoring pathways. The side views of the 3 spokes are shown. The NE surface is indicated by wireframe model (grey). The uncertainty of each Nup localization is indicated by the density of the corresponding color: Y-complex (green), isolated fraction of Nup155 (orange), Nup93-Nup188-Nup155 and Nup205-Nup93-Nup155 complexes (blue), and Nup62-Nup58-Nup54 complex (purple). The posterior model likelihood for each pathway is indicated (see Methods for details). The variations in the structural models for the intermediates at 5 and 6 min after anaphase onset would be due to their lower protein density (lower signal-to-noise ratio) in the electron tomography images.

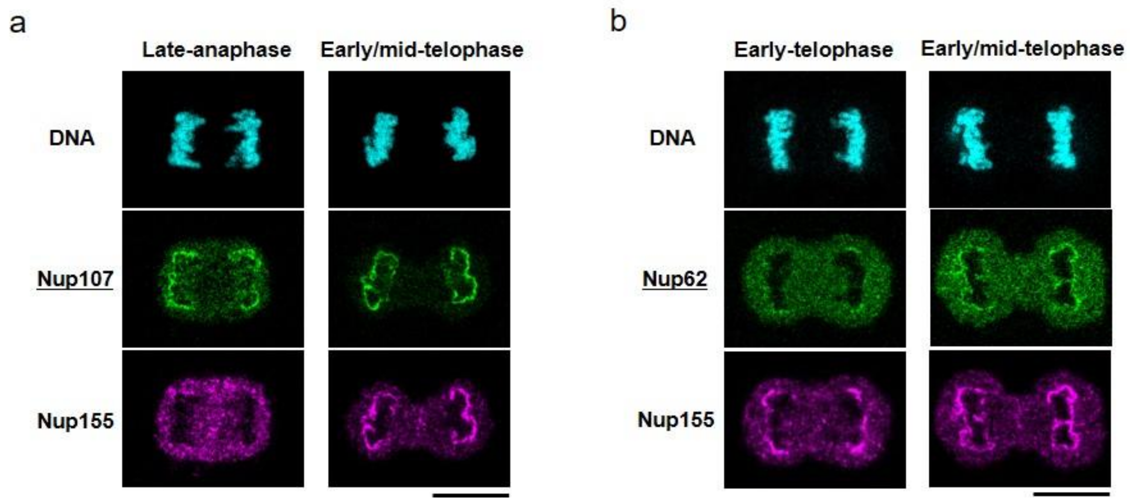


**Extended Data Fig. 8 | Nup155 assembles at a similar timing to Nup62 in postmitotic assembly pathway.**

**a, b**, Immunofluorescence with an anti-Nup155 antibody and a GFP-Nanobody. Cells at anaphase and telophase were imaged by confocal microscopy. mEGFP-Nup107 (**a**) and Nup62-mEGFP (**b**) genome-edited cells were used. The cell nuclei were stained with SiR-DNA. Scale bars, 10 µm.

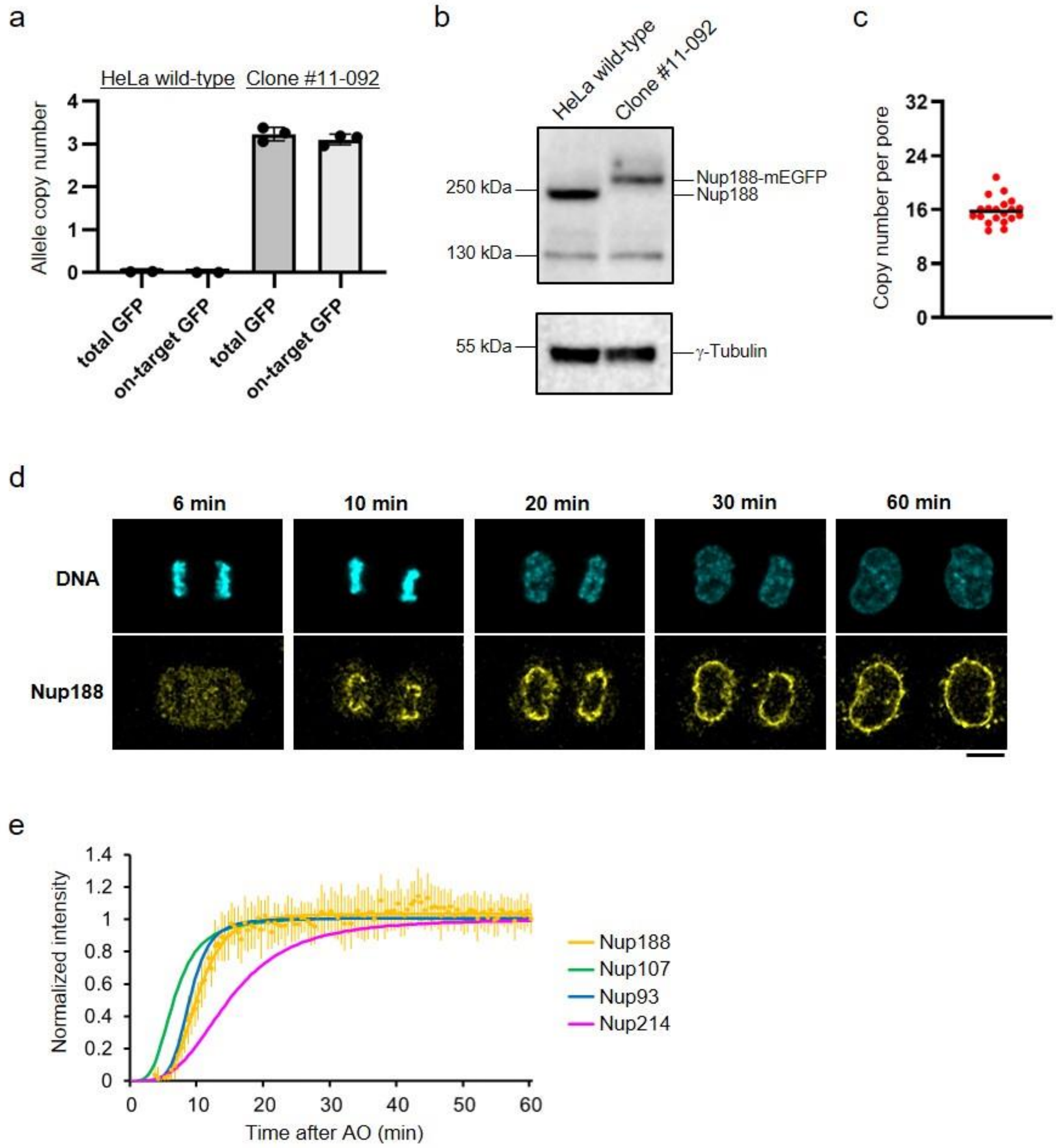


**Extended Data Fig. 9 | Postmitotic assembly kinetics of Nup188 is similar to other inner ring components.**

**a,** Digital PCR analysis of Nup188-mEGFP genome-edited cells for assessing the homozygous tagging. Allele copy numbers of GFP integrations and on-target GFP integrations are quantified. The data are from 2 and 3 measurements for wild-type and Nup188-mEGFP genome-edited HeLa cells. Error bars represent the s.d. of the mean. **b,** Immunoblot analysis of Nup188-mEGFP cells using antibodies against Nup188 and γ-Tubulin. Uncropped immunoblot data are shown in Supplementary Fig. 1. **c,** Calculated copy number of Nup188 per nuclear pore as in Fig. 1. The plot is from 20 cells. The median is depicted as a line. **d,** Time-lapse 3D imaging of Nup188-mEGFP genome-edited cell. The cell nuclei were stained with silicon–rhodamine (SiR)-DNA. Single confocal sections of SiR and GFP channels are shown. Images were filtered with a median filter (kernel size: 0.25 × 0.25 μm). Scale bar, 10 μm. Time after AO is indicated. **e,** Postmitotic assembly kinetics of Nup188 was decomposed as in Extended Data Fig. 4. Dots represent the average and s.d. of measurements from 16 cells. The kinetics of Nup107, Nup93 and Nup214 are also shown for comparison.

| | Interphase | | | Metaphase | |
|---|---|---|---|---|---|
| | Cytoplasm | Nucleoplasm | Nuclear envelope | Cytosol | Nucleoplasm |
| Nup107 | 62 ± 17 (133) | 93 ± 20 (133) | 650 ± 63 (47) | 200 ± 33 (49) | 240 ± 36 (12) |
| Seh1 | 61 ± 16 (31) | 56 ± 14 (32) | 560 ± 61 (37) | 180 ± 26 (20) | 250 ± 28 (12) |
| Nup205 | 15 ± 7.9 (34) | 23 ± 7.7 (34) | 350 ± 54 (41) | 90 ± 14 (21) | 59 ± 5.1 (10) |
| Nup93 | 60 ± 21 (12) | 65 ± 12 (12) | 860 ± 110 (20) | 320 ± 29 (10) | 210 ± 11 (8) |
| Nup62 | 170 ± 52 (28) | 84 ± 27 (28) | 1000 ± 210 (41) | 400 ± 54 (15) | 240 ± 22 (11) |
| Nup214 | 34 ± 12 (16) | 22 ± 2.8 (16) | 450 ± 70 (26) | 120 ± 26 (15) | 68 ± 7.9 (18) |
| Tpr | 6.2 ± 8.3 (12) | 45 ± 18 (12) | 420 ± 63 (55) | 120 ± 20 (10) | 84 ± 19 (9) |
| Nup358 | 13 ± 7.5 (6) | 9.7 ± 3.4 (6) | 390 ± 61 (28) | 88 ± 18 (10) | 65 ± 5.0 (14) |

**Extended Data Table 1 | Concentration of nucleoporins (nM).** Concentration of mEGFP-tagged nucleoporins in interphase and metaphase. Data represent mean ± S.D. from the number of cells indicated in parentheses.
